# PacL-organized membrane-associated effluxosomes coordinate multi-metal resistance in *Mycobacterium tuberculosis*

**DOI:** 10.1101/2025.03.25.645379

**Authors:** Pierre Dupuy, Yves-Marie Boudehen, Marion Faucher, John A. Buglino, Allison Fay, Sylvain Cantaloube, Yasmina Grimoire, Julien Marcoux, Florian Levet, Laetitia Bettarel, Bertille Voisin, Jérôme Rech, Jean-Yves Bouet, Olivier Saurel, Jean-Baptiste Sibarita, Michael Glickman, Claude Gutierrez, Olivier Neyrolles

## Abstract

Metal ion homeostasis is crucial for bacterial pathogens to withstand metal-induced stress during infection. However, the mechanisms underlying bacterial resistance to metal stress remain incompletely understood, particularly how bacteria coordinate responses to simultaneous exposure to multiple metals. Here, we uncover a previously unrecognized mechanism by which *Mycobacterium tuberculosis*, the causative agent of tuberculosis, orchestrates a coordinated response to multi-metal stress. We demonstrate that *M. tuberculosis* assembles dynamic, membrane-associated platforms, organized by PacL proteins, that confer resistance to multiple metals simultaneously. PacL proteins function as scaffolds, clustering multiple P-type ATPase (P-ATPase) pumps, CtpC, CtpG, and CtpV, into functional complexes we term “effluxosomes”. Our findings show that PacL proteins are critical for stabilizing CtpG within membrane-associated clusters, conferring cadmium tolerance, while CtpC serves as a backup, promoting cross-resistance to both zinc and cadmium. Using super-resolution microscopy and single-particle tracking, we elucidate the 3D structure and dynamics of effluxosomes in the mycobacterial membrane. We further demonstrate that conserved residues within the transmembrane domain of PacL proteins are crucial for the assembly of dynamic effluxosomes, which are essential for P-ATPase activity. Additionally, we reveal that PacL1 exhibits metallochaperone activity, binding zinc, cadmium, and copper via a conserved C-terminal motif. Proximity labeling further identifies an extensive PacL1 interaction network, encompassing multiple proteins involved in stress adaptation. Our findings introduce effluxosomes as dynamic, membrane-associated efflux machineries that mediate coordinated multi-metal resistance in *M. tuberculosis*, providing new insights into bacterial metal homeostasis and unveiling potential antimicrobial targets.

## Introduction

Bacteria must tightly regulate metal ion homeostasis to maintain cellular function and adapt to fluctuating environmental conditions. Transition metals are essential for numerous biochemical processes, acting as cofactors for enzymes and as structural components of proteins. However, an imbalance in these metals can be toxic. Our research, along with that of others, has shown that immune cells exploit transition metals to intoxicate bacterial pathogens, and that metal efflux systems contribute to bacterial virulence^1–4^. Specifically, we demonstrated that macrophage phagosomes containing *Mycobacterium tuberculosi*s, the causative agent of tuberculosis, accumulate toxic levels of zinc^2^. This discovery was made together with the identification of an unprecedented mycobacterial zinc detoxification system composed of a membrane P-type ATPase metal efflux pump, CtpC^2^, and a previously uncharacterized protein, PacL1^5^, containing a domain of unknown function DUF1490. CtpC plays a key part in both *M. tuberculosis* zinc tolerance^2^ in culture and in bacterial replication within host cells and tissues^2,6–10^. PacL1, a small membrane-associated protein, is indispensable for CtpC-dependent zinc tolerance^5^. PacL1 directly interacts with CtpC, and both proteins colocalize in dynamic membrane patches. Acting as a chaperone, PacL1 plays a critical role in stabilizing CtpC and also exhibits putative metallochaperone activity, which is mediated by a C-terminal metal-binding motif essential for zinc tolerance under high-zinc conditions.

Intriguingly, *M. tuberculosis* possesses two homologs of PacL1, namely PacL2 and PacL3^5^, whose biological functions remain unknown. All PacL proteins share common structural features, including an N-terminal transmembrane (TM) α-helix, a cytoplasmic domain containing Ala/Glu (AE) repeats, and a C-terminal intrinsically disordered region (IDR). PacL2 and PacL3 are encoded within an operon alongside two putative P-type ATPases, CtpG and CtpV, as well as two transcriptional regulators, CmtR and CsoR. These regulators induce operon expression in response to cadmium^11,12^ and copper^13^, respectively. However, conflicting studies suggest that CtpG contributes to tolerance against zinc^9^, cadmium^14^, and copper^14^, while CtpV appears to be primarily associated with copper tolerance^10^. Notably, both CtpG and CtpV, like CtpC, are induced during macrophage infection^15^ and contribute to bacterial growth within the host^9,10^. Whether the enzymatic activities of CtpG and CtpV depend on PacL proteins remains unknown. Interestingly, our previous research demonstrated that when PacL1, PacL2, and PacL3 are co-expressed as an artificial operon under a constitutive promoter in the *M. tuberculosis* non-pathogenic relative *Mycobacterium smegmatis*, they colocalize within shared membrane clusters^5^. This suggests a close functional collaboration between PacL proteins and various P-ATPase pumps in *M. tuberculosis*.

Here, we reveal a previously unrecognized interplay between PacL proteins and diverse P-type ATPase pumps in *M. tuberculosis*, uncovering the existence of atypical multi-metal efflux machineries. By dissecting the functional roles of PacL2 and CtpG, we establish their specific involvement in cadmium tolerance and demonstrate that PacL2 is essential for both the stability and membrane-associated localization of CtpG. Furthermore, our findings highlight a broader interaction network in which PacL1, PacL2, and PacL3 physically associate with each other, as well as with CtpC, CtpG, and CtpV, collectively contributing to multi-metal resistance. Using super-resolution photoactivated localization microscopy (PALM), we characterize these multiprotein membrane clusters and identify conserved residues critical for their assembly. Finally, using proximity-ligation assay, we map the PacL1 interaction network, uncovering the potential involvement of dozens of additional proteins within these clusters. These findings not only deepen our understanding of metal homeostasis in *M. tuberculosis* but also suggest that similar dynamic membrane multiprotein machineries, involving PacL-Ctp pairs found across a variety of bacterial species and genera, may represent a conserved strategy for adapting to metal stress, offering new perspectives on microbial resilience and potential targets for antimicrobial intervention.

## Results

### PacL/Ctp systems mediate dual tolerance to zinc and cadmium

To investigate the role of the three PacL/Ctp systems in *M. tuberculosis* metal tolerance, we expressed the *pacL1-ctpC*, *cmtR-pacL2-ctpG*, and *csoR-pacL3-ctpV* operons, along with their native promoters, in *M. smegmatis* (**Fig. 1a**). Using a disc diffusion assay, we observed that the expression of *pacL1-ctpC* and *pacL2-ctpG* conferred increased tolerance to zinc (**Fig. S1a**) and cadmium (**Fig. S1b**), respectively, but not to other tested metals, including copper (**Fig. S1c**), nickel (**Fig. S1d**), manganese (**Fig. S1e**), or iron (**Fig. S1f**). Mutation of the conserved APC motif in the P-ATPase domain of CtpG (APC→AAA), which is predicted to be essential for metal binding and transport^3^, abolished the enhanced cadmium tolerance conferred by the expression of the *ctpG* operon (**Fig. S1b**).

**Figure 1.**
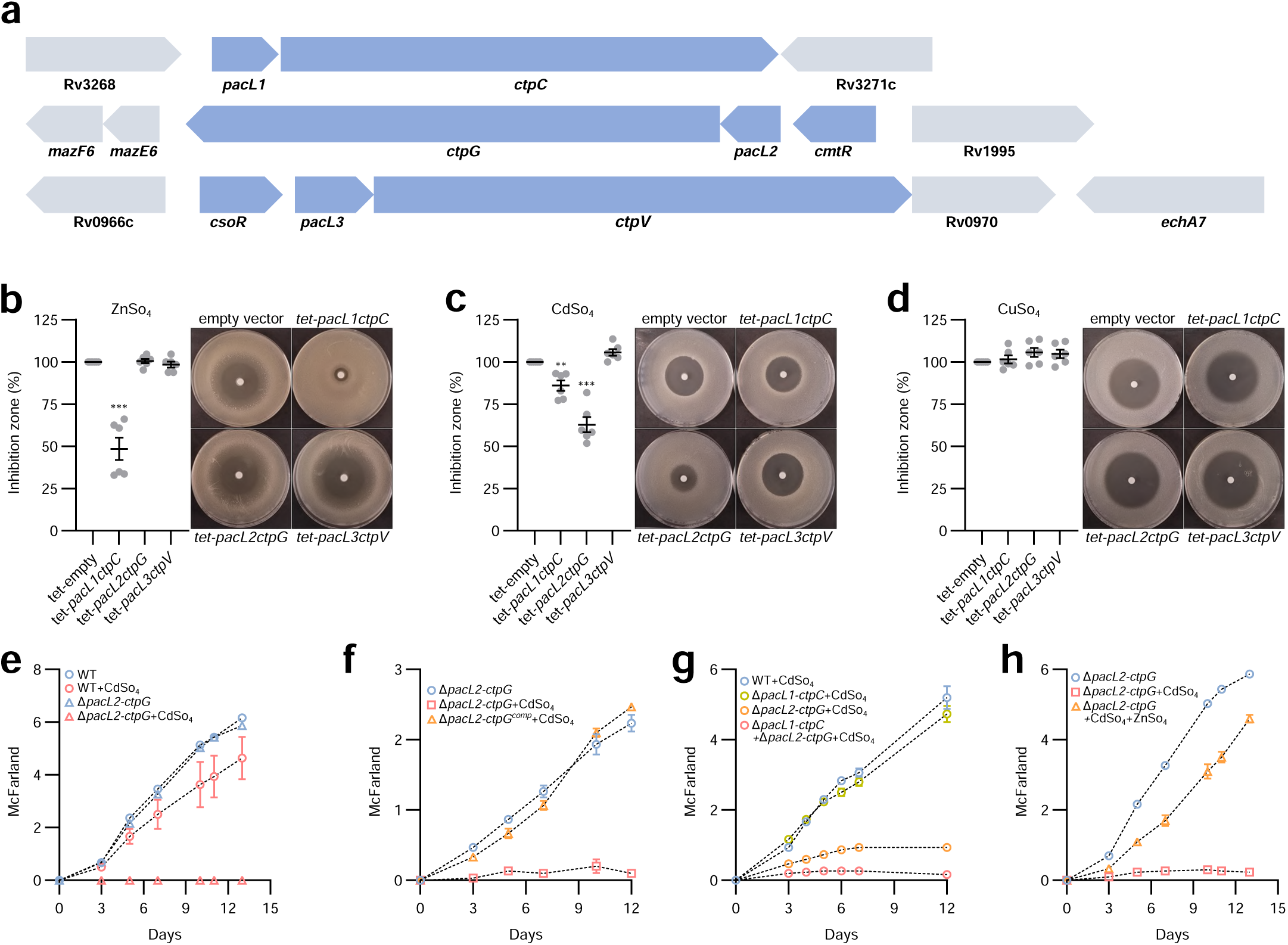
CtpC and CtpG collaborate to confer cadmium tolerance in *M. tuberculosis*. (**a**) Genomic organization of *pacL*, *ctp*, and adjacent genes in the *M. tuberculosis* chromosome. (**b**–**d)** Sensitivity of the indicated *M. smegmatis* strains to the specified metals, assessed by disk diffusion assay in the presence of an inducer (ATC). Results represent the diameter of the inhibition zone, expressed relative to the empty vector condition. *tet*: ATC-inducible promoter. Results are presented as means (±SEM) from biological replicates, represented by gray dots. Asterisks above the means indicate statistically significant differences compared to the reference strain (empty vector) (**P < 0.01; ***P < 0.001). (**e**–**h**) Bacterial growth of the indicated *M. tuberculosis* strains in the absence or presence of the specified metals (20 µM CdSo_4_ and 50 µM ZnSO_4_), evaluated by turbidity measurement (reported in McFarland units). Data show mean ± SEM of a biological triplicate.

However, the expression of the *ctpV* operon did not affect tolerance to any of the tested metals (**Fig. S1a, b, c, d, e,** and **f**). As this result contradicts a previous study suggesting that CtpV contributes to copper tolerance^10^, we additionally expressed the *csoR-pacL3-ctpV* operon together with the downstream *Rv0970* gene (**Fig. 1a**) to assess whether this gene plays a role in CtpV function. Nevertheless, we observed no effect of the full operon expression on copper tolerance in *M. smegmatis* when tested with either Cu^2+^ (**Fig. S1g**) or Cu^+^ (**Fig. S1h**).

Previous studies have shown that *ctpC*, *ctpG*, and *ctpV* are induced by zinc^2^, cadmium^11^, and copper^10^, respectively. To confirm that the observed metal specificity of the PacL1/CtpC and PacL2/CtpG systems arises from their intrinsic enzymatic activities rather than differential expression triggered by specific stress conditions, we expressed each PacL/Ctp pair under an anhydrotetracycline (ATC)-inducible (*tet*) promoter^16^. In the absence of the inducer, no differences in metal tolerance were detected between strains (**Fig. S2a**). Constitutive expression of *pacL3-ctpV* did not confer increased tolerance to any tested metal, whereas *pacL2-ctpG* expression specifically enhanced cadmium tolerance (**Fig. 1b, 1c, 1d**, and **S2b**). Interestingly, constitutive expression of *pacL1-ctpC* conferred increased tolerance to both zinc and cadmium (**Fig. 1b** and **1c**), suggesting that, when sufficiently expressed, this system can facilitate the efflux of both metals.

To validate the role of PacL1/CtpC and PacL2/CtpG systems in cadmium tolerance in *M. tuberculosis*, we generated deletion mutants for each system individually and in combination (Δ*pacL1ctpC*, Δ*pacL2ctpG, and* Δ*pacL1ctpC*Δ*pacL2ctpG*). Deletion of *pacL2* and *ctpG* resulted in increased sensitivity to cadmium (**Fig. 1e**), whereas ectopic expression of these genes in the mutant background fully restored cadmium tolerance (**Fig. 1f**). Notably, the Δ*pacL1ctpC*-*pacL2ctpG* double mutant exhibited even greater sensitivity to cadmium than the Δ*pacL2ctpG* single mutant (**Fig. 1g**), suggesting that the PacL1/CtpC system serves as a backup mechanism for PacL2/CtpG in mediating cadmium tolerance. Given that *ctpC* expression is induced by zinc in *M. tuberculosis*, we further investigated whether zinc supplementation could influence cadmium tolerance in the Δ*pacL2ctpG* mutant background. Interestingly, zinc enhanced cadmium tolerance in *M. tuberculosis* (**Fig. 1h**), revealing a cross-resistance mechanism between zinc and cadmium in this pathogen.

### PacL2 is essential for stabilizing CtpG within membrane-associated clusters

To elucidate the role of PacL2 in CtpG-dependent cadmium tolerance in *M. tuberculosis*, we expressed *ctpG* in *M. smegmatis*, either alone or in combination with *pacL2*. While co-expression of *pacL2* and *ctpG* conferred significant cadmium tolerance, expression of *pacL2* alone had no effect, and *CtpG* expression alone provided only weak tolerance (**Fig. 2a**). To further validate the role of PacL2 in *M. tuberculosis*, we generated an in-frame deletion mutant of *pacL2*. Deletion of *pacL2* led to increased sensitivity to cadmium (**Fig. 2b**), which was fully reversed by ectopic expression of *pacL2* in the mutant background (**Fig. 2c**), confirming its essential role in cadmium tolerance.

**Figure 2.**
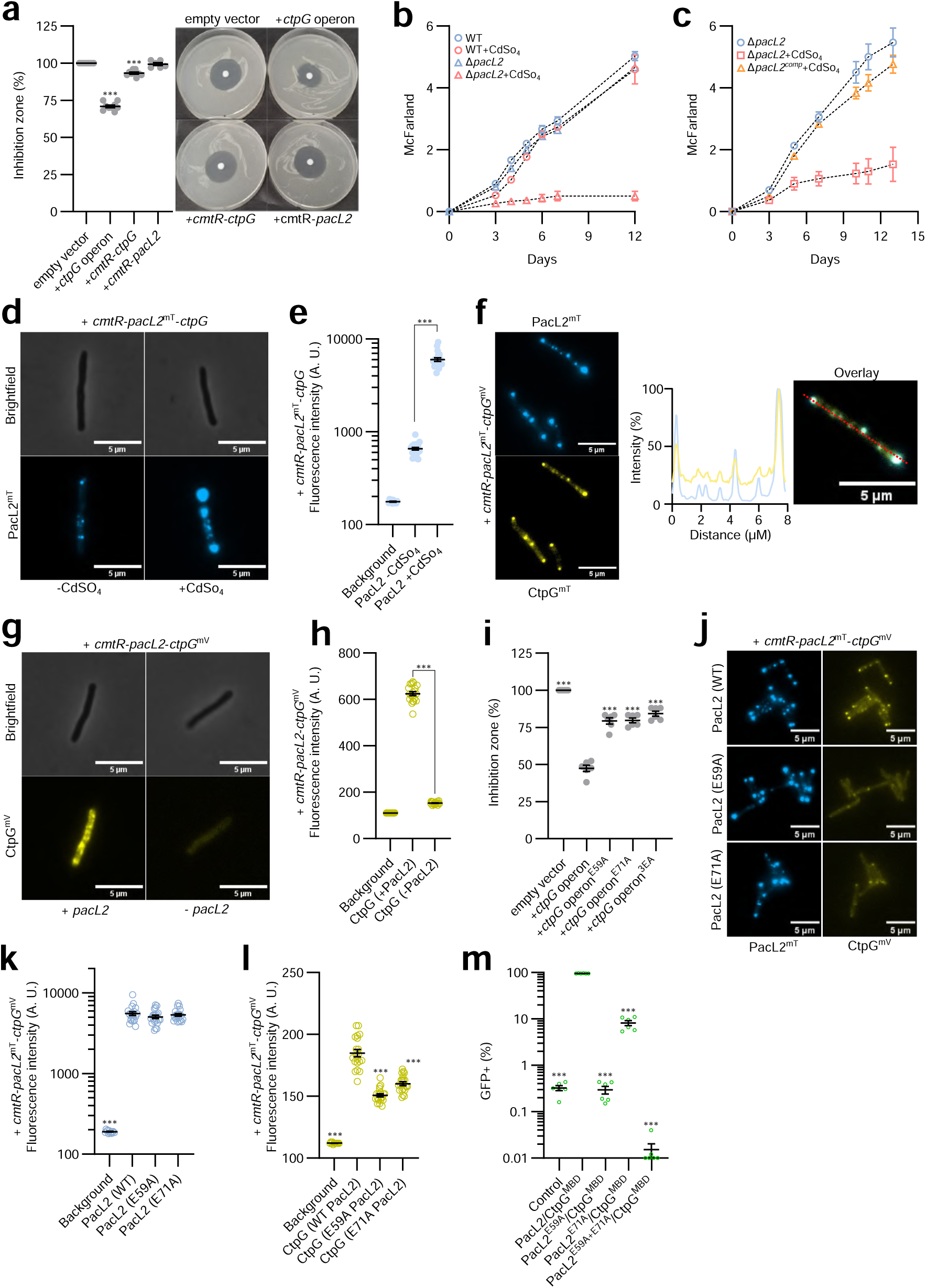
PacL2 is essential for stabilizing CtpG within membrane clusters. (**a** and **i**) Sensitivity of the indicated *M. smegmatis* strains to CdSo_4_, assessed by disk diffusion assay. Results represent the diameter of the inhibition zone, expressed relative to the empty vector condition. Data are presented as means (±SEM) from biological replicates, represented by gray dots. *ctpG* operon: *cmtR-pacL2-ctpG*. (**b, c**) Bacterial growth of the indicated *M. tuberculosis* strains in the absence or presence of 20 µM CdSo_4_, evaluated by turbidity measurement (reported in McFarland units). Data represent mean ± SEM from biological triplicates. (**d, f, g**, and **j**) Epifluorescence microscopy images of indicated *M. smegmatis* strains expressing PacL2-mTurquoise and/or CtpG-mVenus cultivated with 100 nM CdSO_4_. (**e, h, k**, and **l**) Fluorescence intensity measurements in grayscale of (blue) PacL2-mTurquoise or (yellow) CtpG-mVenus proteins expressed in the indicated strains. Results are presented as means (±SEM) from individual cells represented by circles. The right panel in (**f**) shows co-localization of mTurquoise and mVenus fluorescent signals along the red line in the overlay image. (**m**) Bipartite split-GFP assay in *M. smegmatis* expressing a PacL1-GFP11 fusion protein along with GFP1-10 (control) or the indicated proteins fused to GFP11 (first protein) or GFP1-10 (second protein). CtpG^MBD^: N-terminal domain of CtpG encoding the predicted metal binding domain of the P-ATPase. Fluorescence was recorded by flow cytometry and expressed as the percentage of GFP-positive cells. Data are presented as means (±SEM) from biological replicates, represented by green dots. Asterisks indicate statistically significant differences compared to the reference condition (empty vector (**a** and **i**), background (**k**), CtpG (WT PacL2) (**l**), or PacL2/CtpG^MBD^ (**m**)) or between conditions linked by a black line (**e, h**) (***P < 0.001).

To determine the cellular localization of PacL2 and CtpG, we engineered PacL2-mTurquoise and CtpG-mVenus fusion proteins. Using epifluorescence microscopy, we observed that PacL2 localized in distinct clusters, which became more intense and abundant upon cadmium exposure (**Fig. 2d** and **e**). Under the same conditions, CtpG also formed clusters that co-localized with PacL2 (**Fig. 2f**). Interestingly, in the absence of PacL2, CtpG clusters were undetectable (**Fig. 2g** and **h**), indicating that PacL2 is essential for stabilizing CtpG and facilitating its proper localization within membrane-associated clusters.

Our previous study demonstrated that AE repeats located in the cytoplasmic domain of PacL1 are essential for both the PacL1/CtpC interaction and the stabilization of CtpC. Given that similar AE repeats are also present in the PacL2 sequence (**Fig. S3**), we investigated their role in CtpG stability. Substitutions E59A or E71A, or the triple substitution E55A, A59A, and E71A abolished the resistance to Cd when the mutated operon was expressed in *M. smegmatis* (**Fig. 2i**). In addition, although E59A or E71A substitutions did not prevent the formation of PacL2-mTurquoise clusters, they resulted in destabilization of CtpG-mVenus (**Fig. 2j, k**, and **l**). Finally, bipartite split-GFP experiments demonstrated that PacL2 interacts with the predicted metal binding domain (MBD) of CtpG and that EA substitutions in PacL2 reduced this interaction (**Fig. 2m**). Altogether, these data demonstrate that the acidic residues present in conserved AE motifs on the PacL2 cytoplasmic domain are essential for its protein chaperone activity. Together, these results suggest that PacL2 functions as a scaffold protein, facilitating the recruitment of CtpG into membrane-associated clusters and protecting it from degradation.

### PacL2 proteins clustering and dynamics

Analysis of the dynamics of the PacL2-CtpG clusters revealed two distinct types of structures: large, immobile clusters frequently localized at the cell poles, and smaller, mobile clusters that moved dynamically throughout the cell (<u>Movie 1</u>). The smaller clusters appeared to preferentially localize in specific regions where they interact with one another. To further characterize the spatial organization of PacL2 and CtpG, we generated PacL2-mEos and CtpG-mEos fusion proteins for super-resolution microscopy. Using 3D PALM, we constructed the tridimensional nanoscale organization of PacL2 and CtpG within individual *M. smegmatis* cells (**Fig. 3a, 3b**, and <u>Movie 2</u>). Using the point cloud visualization and segmentation software PoCA^17^ we quantified the number of clusters (**Fig. 3c**), their volumes (**Fig. 3d**), and the proportion of PacL2 molecules localized within clusters (**Fig. 3e**). Our analysis revealed that approximately 60% of cellular PacL2 was organized into an average of 12.5 membrane clusters per cell (**Fig. 3c** and **e**), with cluster sizes ranging from 10^5^ nm³ to 10^8^ nm³ (**Fig. 3d**). CtpG formed nearly as many clusters per cell as PacL2 (**Fig. 3c**). However, the CtpG clusters were smaller (**Fig. 3d**) and less dense, with fewer than 20% of CtpG molecules localized within clusters (**Fig. 3e**).

**Figure 3.**
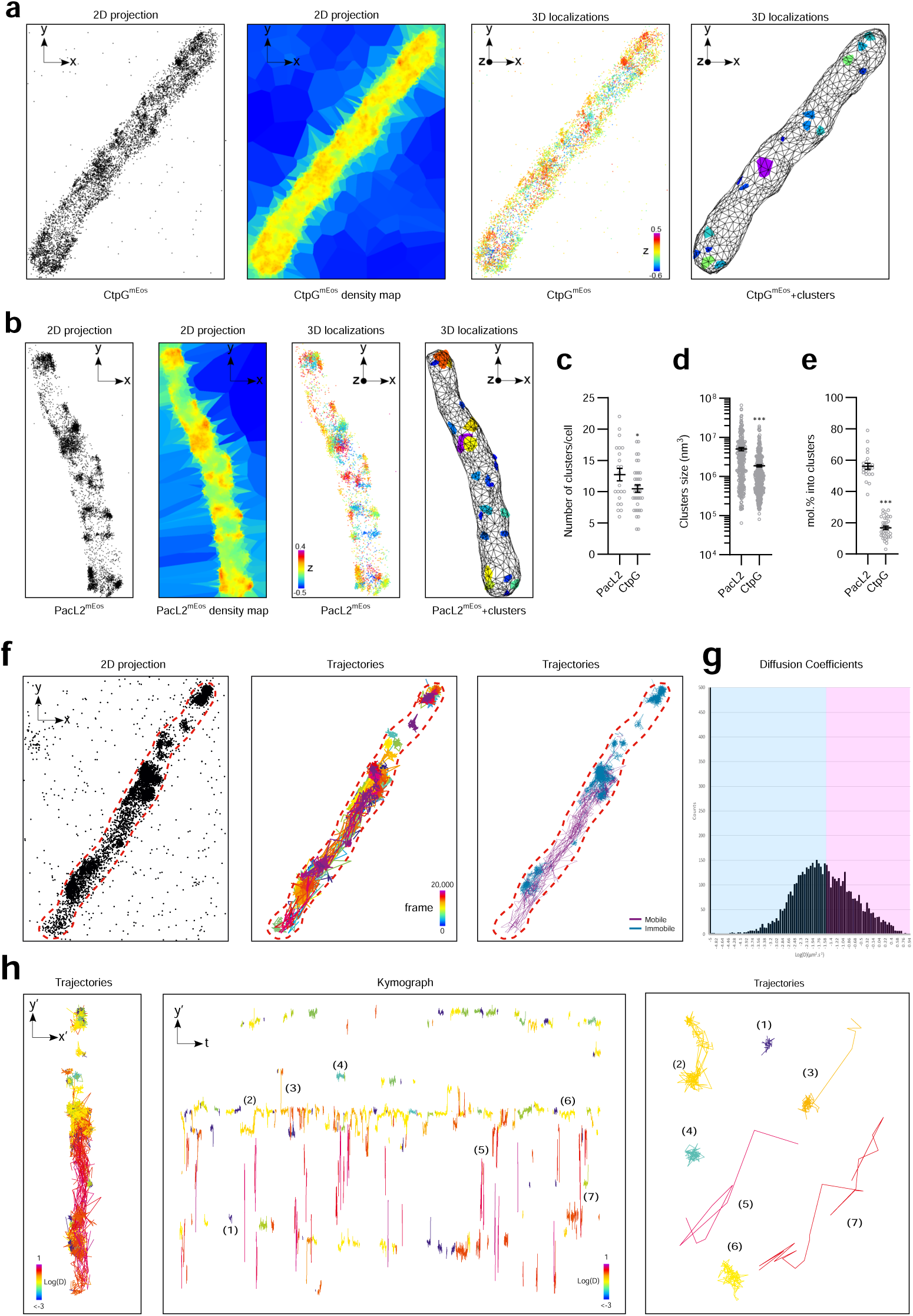
**PacL2 proteins clustering and dynamics**. (**a** and **b**) Representative 3D super-resolution reconstructions of PacL2-mEos or CtpG-mEos proteins in *M. smegmatis* cells using 3D-PALM. From left to right: localizations, density map, 3D localizations, and cluster segmentation using the PoCA software. Quantification of (**c**) the number of PacL2 or CtpG clusters per cell, (**d**) cluster sizes, and (**e**) proportion of clustered molecules per cell. Results are presented as means (±SEM) from individual cells (**c, e**) or clusters (**d**), represented by gray dots. (**f**) Representative super-resolution reconstructions of PacL2-mEos protein acquired by sptPALM. From left to right: localizations, trajectories with lengths >6 time points, overlay between slow (cyan) and fast (magenta) molecules. (**g**) Histogram of diffusion coefficients computed from 24 cells. (**h**) Trajectories and kymograph analysis along the long axis of the cell, color-coded with their diffusion coefficients. Examples of trajectories show immobilized, freely diffusive, and highly mobile molecules.

We then monitored the mobility of PacL protein using single protein tracking combined with PALM (sptPALM). Analyzing the localization and dynamics of single PacL molecules over 2.5 minutes at 33 frames per seconds, we could retrieve the nanoscale organization of PacL in nanoclusters of different sizes distributed along the bacteria, as well as its dynamic behavior (**Fig. 3f-h**). Diffusion analysis computed from the mean square displacements (MSD) of individual single molecule trajectories (7,453 trajectories longer than 6 time points extracted from 24 isolated cells) revealed a broad distribution of mobility integrating mobile and immobile molecules, without a clear cut-off between the two populations (**Fig. 3g**). To further characterize and map mobile and immobile molecules, we applied an experimental threshold of 0.03 µm²·s[¹ (**Fig. 3g**). Our analysis revealed that 70% of the molecules were immobile, while 30% were classified as mobile. These two populations were spatially mapped within the bacteria, clearly distinguishing confined molecules within clusters from mobile molecules moving between them (**Fig. 3f**). Using kymograph analysis to investigate the spatio-temporal distribution of individual PacL molecules along the bacterial long axis (**Fig. 3h**), we identified three distinct behaviors: (i) confined molecules forming large, stable clusters visible throughout the acquisition; (ii) transiently confined molecules, likely representing small, slowly mobile clusters, characterized by brief immobilizations; and (iii) highly mobile molecules traveling along the bacteria, depicted as long red traces. Only rare events of state transitions from immobile to mobile were observed.

### PacL proteins function as hub molecules that organize multiple P-ATPase pumps into shared membrane-associated clusters

Our previous study demonstrated that PacL1 colocalizes with PacL2 and PacL3 within shared membrane clusters when these genes are artificially co-expressed as an operon under the control of a constitutive promoter in *M. smegmatis*^5^. This observation raises the question of whether these proteins physically interact. To investigate this, we performed bipartite split-GFP experiments and found that PacL1 interacts not only with itself but also with both PacL2 and PacL3, with PacL2 exhibiting a similar interaction pattern (**Fig. 4a**). As a negative control, we observed a four-fold lower GFP signal when testing the interaction between PacL1 and the mycobacterial flotillin-like Rv1488, a protein known to promote membrane clusters distinct from PacL-dependent platforms^5^.

**Figure 4.**
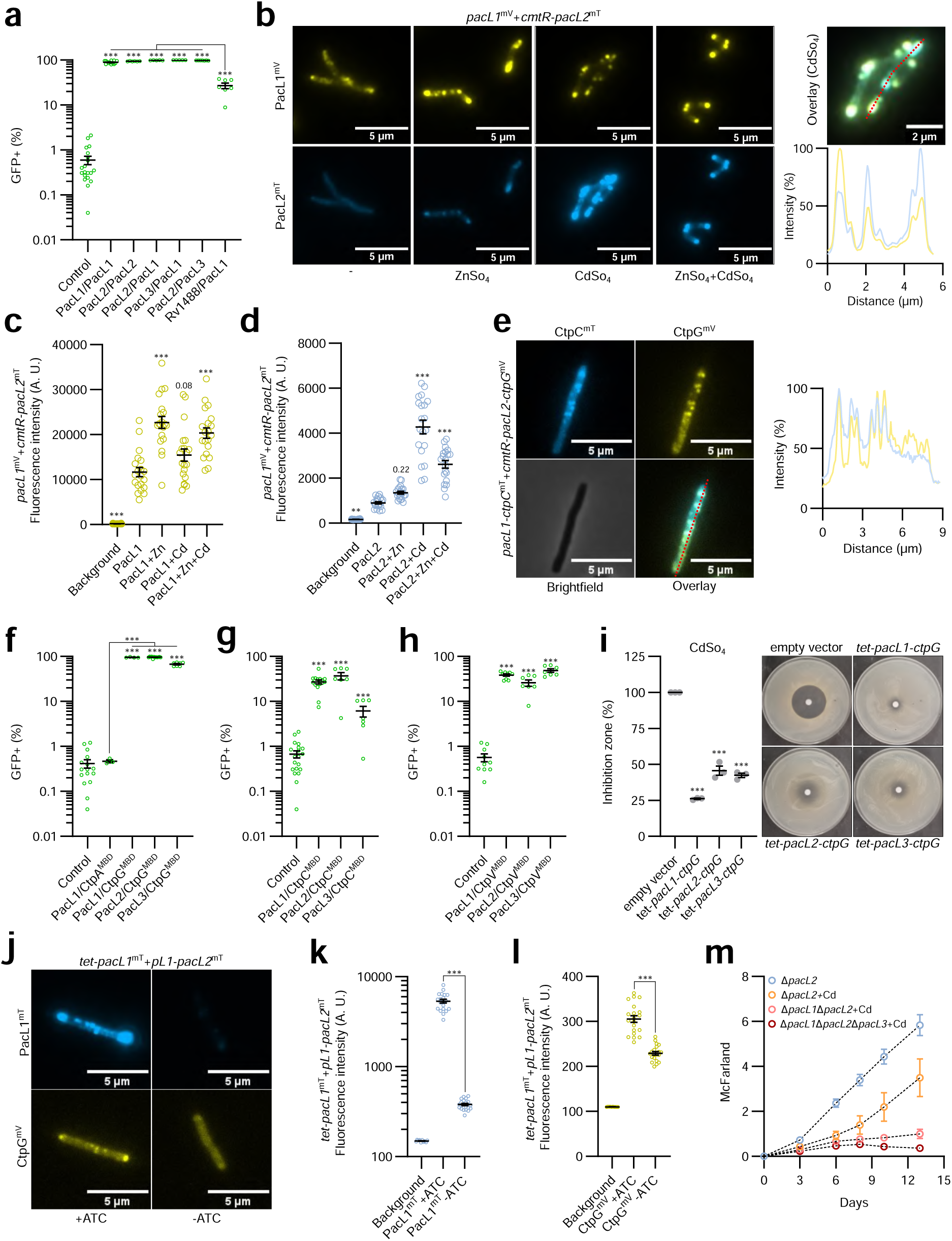
PacL proteins collaborate to assemble multi-metal efflux machineries. (**a, f**–**h**) Bipartite split-GFP assay in *M. smegmatis* expressing a PacL1-GFP11 fusion protein along with GFP1-10 (control) or the indicated proteins fused to GFP11 (first protein) or GFP1-10 (second protein). Fluorescence was recorded by flow cytometry and expressed as the percentage of GFP-positive cells. Data are presented as means (±SEM) from biological replicates, represented by green dots. (**b, j**) Epifluorescence microscopy images of (**b**) fixed *M. tuberculosis* or (**e** and **j**) live *M. smegmatis* expressing the indicated fusion proteins. Bacteria were cultured in the absence or presence of (**b**) the indicated metals (100 µM ZnSO_4_ and 10 µM CdSO_4_), (**e**) 100 nM CdSO_4_ or (**j**) 50 nM anhydrotetracycline (ATC). *tet*-: ATC inducible promoter, *pL1*: pacL1 promoter. The right panels in (**b** and **e**) show co-localization of mTurquoise and mVenus fluorescent signals along the red line in the overlay image. (**c, d, k, l**) Fluorescence intensity measurements of the indicated strains and conditions in grayscale. Results are presented as means (±SEM) from individual cells, represented by yellow or blue dots. (**i**) Sensitivity of the indicated *M. smegmatis* strains to cadmium, assessed by disk diffusion assay in the presence of the inducer (ATC). Results represent the diameter of the inhibition zone, expressed relative to the empty vector condition. *tet*: ATC-inducible promoter. Data are presented as means (±SEM) from biological replicates, represented by gray dots. (**m**) Bacterial growth of the indicated *M. tuberculosis* strains in the absence or presence of 10 µM CdSO_4_, evaluated by turbidity measurement (reported in McFarland units). Data represent mean ± SEM from biological triplicates. Asterisks indicate statistically significant differences compared to the reference condition (Control (**a, f, g, h**), PacL1 (**c**), PacL2 (**d**), empty vector (**i**) or between conditions linked by a black line (**f, k, l**) (*P < 0.05; **P < 0.01; ***P < 0.001).

To determine whether PacL/PacL colocalization occurs in *M. tuberculosis* under more physiological conditions, we independently expressed PacL1-mVenus and PacL2-mTurquoise fusion proteins under the control of their respective native promoters, either in the absence of metals or in the presence of zinc, cadmium, or both. PacL1 clusters were consistently detected in *M. tuberculosis* membranes under all four conditions (**Fig. 4b** and **c**), with an increase in fluorescent intensity upon metal exposure, particularly in response to zinc, which induced the highest signal (**Fig. 4c**). In contrast, in the absence of metals, PacL2-mTurquoise fluorescence was very weak and displayed a homogeneous distribution within *M. tuberculosis* cells (**Fig. 4b** and **d**). Notably, exposure to both zinc and cadmium triggered the formation of membrane-associated PacL2 clusters, with cadmium eliciting the strongest increase in fluorescence signal (**Fig. 4b** and **d**). Under metal stress conditions, we showed the colocalization of PacL1 and PacL2 within the *M. tuberculosis* membrane (**Fig. 4b**). These results suggest that environmental conditions modulate the composition of PacL-dependent platforms, with zinc preferentially promoting PacL1-enriched clusters and cadmium favoring the formation of PacL2-enriched clusters.

Next, we explored whether the clustering of PacL1 and PacL2 also facilitates the co-clustering of CtpC and CtpG. To test this hypothesis, we expressed PacL1 and CtpC fused to mTurquoise, alongside PacL2 and CtpG fused to mVenus, in *M. smegmatis* under the control of their respective native promoters. Remarkably, we observed that both P-type ATPase pumps colocalized within shared clusters (**Fig. 4e**), revealing the presence of previously unrecognized multi-metal efflux machineries in mycobacteria.

### Redundant chaperone functions of PacL proteins in stabilizing P-type ATPase pumps

Our previous study revealed that PacL1 physically interacts with the MBD of CtpC^5^. Here, we used the bipartite split-GFP assay to investigate potential cross-interactions between PacL proteins and the MBDs of CtpC, CtpG, and CtpV. Interestingly, we found that the MBDs of CtpG and CtpV interact with PacL1, PacL2, and PacL3, whereas the MBD of CtpC interacts with PacL1 and PacL2, but less efficiently with PacL3 (**Fig. 4f, g,** and **h**). In contrast, PacL1 does not interact with the MBD of CtpA (**Fig. 4f**), demonstrating that not all P-type ATPases of *M. tuberculosis* can be partners of PacL1. To assess whether PacL1 can substitute for PacL2 or PacL3 in stabilizing CtpG, we generated *M. smegmatis* strains expressing CtpG in combination with PacL1, PacL2, or PacL3 under the control of a *tet* promoter. Disc diffusion assays revealed that all three PacL proteins conferred CtpG-dependent cadmium tolerance, with PacL1 providing the highest level of resistance (**Fig. 4i**). To confirm the role of PacL1 in stabilizing CtpG within membrane clusters, we co-expressed a CtpG-mVenus fusion protein under a constitutive promoter along with a PacL1-mTurquoise fusion protein under the control of a *tet* promoter. In the presence of the inducer, PacL1 and CtpG formed fluorescent foci that colocalized in the membrane of *M. smegmatis* (**Fig. 4j**). In the absence of the induction, the fluorescence intensity of both PacL1 and CtpG decreased significantly, indicating that PacL1 contributes to CtpG stability (**Fig. 4j, k** and **l**). Finally, to investigate the contribution of PacL proteins to cadmium tolerance in *M. tuberculosis*, we constructed multiple *pacL* knockout mutants. The *pacL1 pacL2* double mutant exhibited greater sensitivity to cadmium than the *pacL1* single mutant (**Fig. 4m**), demonstrating that at least two PacL proteins function cooperatively to confer cadmium tolerance in this pathogen.

### Distinct roles of PacL proteins in metallochaperone activity

PacL1 contains a C-terminal metal-binding motif, D^87^LHDHDH^93^ (**Fig. S3**) that binds zinc at a 1:1 molar ratio and is crucial for resistance to high zinc concentrations in *M. tuberculosis*, supporting its role as a metallochaperone^5^. The clustering of multiple P-type ATPase pumps within shared membrane domains raises the question of whether the metallochaperone function of PacL1 extends beyond zinc binding. To test this hypothesis, we employed native mass spectrometry to assess the relative binding affinity of the purified cytoplasmic domain of PacL1 to various metals. Interestingly, PacL1 was capable of binding cadmium (**Fig. 5a** and **c**), zinc (**Fig. 5d**), and copper (**Fig. 5e**) but not manganese (**Fig. 5f**).

**Figure 5.**
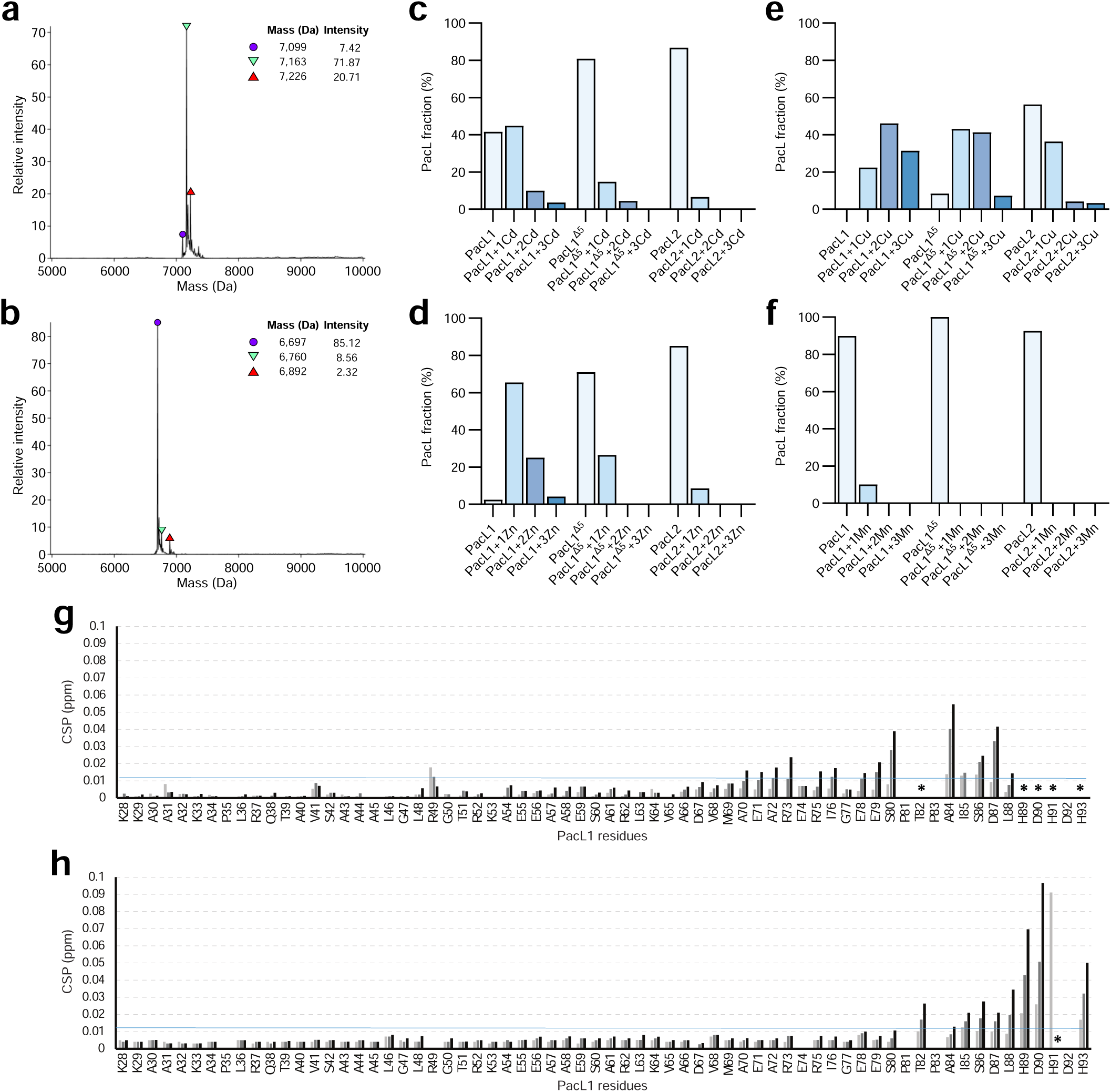
Distinct roles of PacL proteins in metallochaperone activity. (**a, b**) Deconvoluted mass spectra obtained by native MS and showing cadmium binding to (**a**) PacL1 but not to (**b**) PacL1Δ5. (**c, d, e, f**) Proportion of the indicated purified proteins bound to specific numbers of specified metals atoms, assessed by native mass spectrometry. (**g, h**) Residue-specific perturbations induced by (**g**) zinc and (**h**) cadmium on PacL1. Summary of chemical shift perturbations of PacL1 due to zinc and cadmium interaction at 0.4 (light grey), 1.0 (grey) and 2 (black) equivalent of zinc and cadmium. Black stars indicated peak broadening below the detection due to chemical exchange. PacL1: soluble domain of PacL1, PacL1Δ5: soluble domain of PacL1 deleted from its C-terminal metal-binding motif, and PacL2: soluble domain of PacL2.

We then performed NMR spectroscopy experiments to identify the specific residues in PacL1 responsible for zinc and cadmium binding. Chemical shift perturbation (e.g. CSP) of amide bonds showed that the cadmium binding domain clearly overlaps with those involved in zinc coordination, from H_89_ to H_93_ (**Fig. 5g** and **h** and **S4**). Detailed analysis of the 2D ^1^H-^15^N HSQC spectra for increasing concentrations of zinc and cadmium revealed a slightly different binding mode of cadmium compared to zinc, as the amide signals in the presence of cadmium are less prone to signal broadening due to intermediate chemical exchange (**Fig. S4**). Except H_91_D_92_ which are still broadened, the amide peaks involved in the cadmium interaction are shifted to the fast exchange regime, in line with a lower binding affinity for cadmium and also partially due to different conformations of the bound form.

Deletion of the C-terminal metal-binding motif (PacL1^Δ^^5^) significantly reduced its metal-binding capacity (**Fig. 5b, c, d** and **e**), confirming the critical role of this motif in metal coordination. To evaluate the metal-binding ability of PacL2, we conducted a similar analysis using its purified cytoplasmic domain. PacL2 bound copper, albeit with lower affinity than PacL1 (**Fig. 5e**), but displayed no detectable binding to zinc (**Fig. 5d**), cadmium (**Fig. 5c**), or manganese (**Fig. 5f**). Similarly, NMR spectroscopy also confirmed the absence of binding of zinc and cadmium to PacL2 (data not shown).

### The cytoplasmic domain of PacL proteins is not involved in cluster formation

The mechanism by which PacL proteins form membrane clusters remains unclear. To investigate whether the cytoplasmic domain of PacL proteins contributes to PacL/PacL interactions, we constructed PacL2 deletion mutants lacking segments of its cytoplasmic domain containing these repeats (PacL2^Δ55–84^ and PacL2^Δ31–84^) (**Fig. 6a**).

**Figure 6.**
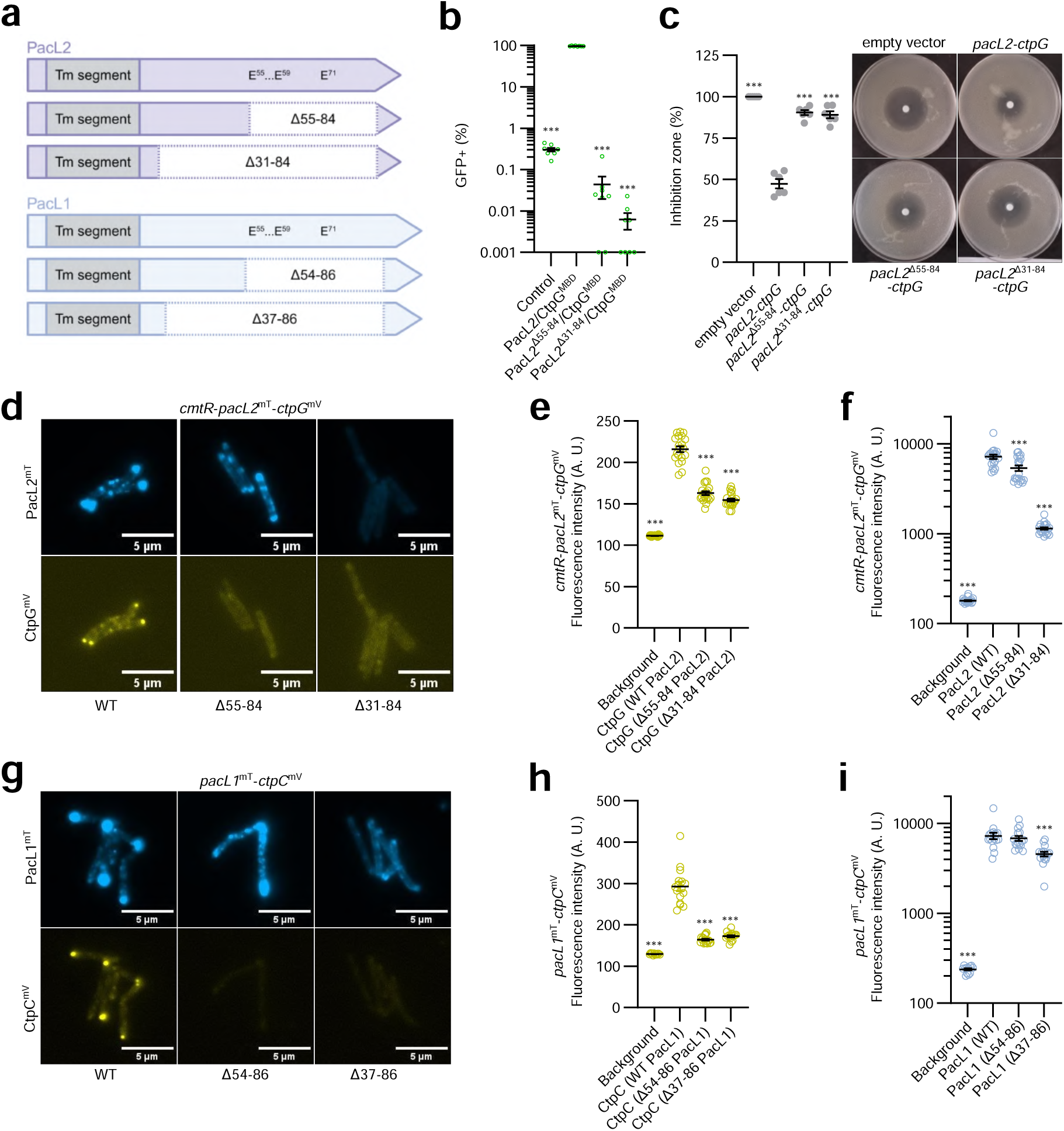
The cytoplasmic domain of PacL proteins is not essential for PacL clustering. (**a**) Schematic representation of PacL1 and PacL2 proteins with deletions in the indicated segments of their cytoplasmic domains. Glutamic acids involved in the PacL/Ctp interaction are marked. (**b**) Bipartite split-GFP assay in *M. smegmatis* expressing a PacL1-GFP11 fusion protein along with GFP1-10 (control) or the indicated proteins fused to GFP11 (first protein) or GFP1-10 (second protein). Fluorescence was recorded by flow cytometry and expressed as the percentage of GFP-positive cells. Data are presented as means (±SEM) from biological replicates, represented by green dots. (**c**) Sensitivity of the indicated *M. smegmatis* strains to CdSo_4_, assessed by disk diffusion assay. Results represent the diameter of the inhibition zone, expressed relative to the empty vector condition. Data are presented as means (±SEM) from biological replicates, represented by gray dots. (**d, g**) Epifluorescence microscopy images of live *M. smegmatis* cells expressing the indicated fusion proteins under their respective native promoters. Cells imaged in (**d**) were cultivated in 7H9 medium supplemented with 100 nM CdSo4. (**e, f, h, i**) Fluorescence intensity measurements of the indicated strains in grayscale. Results are presented as means (±SEM) from individual cells, represented by yellow or blue dots. Asterisks indicate statistically significant differences compared to the reference condition (PacL2*/*CtpG (**b**), *pacL1-ctpG* (**c**), WT PacL1 or PacL2 (**e, f, h, i**)) (*P < 0.05; **P < 0.01; ***P < 0.001).

As expected, these mutants, which harbor a deletion of AE repeats, exhibited impaired interaction with CtpG (**Fig. 6b**), reduced cadmium tolerance (**Fig. 6c**), and decreased CtpG stability, as assessed by fluorescence microscopy (**Fig. 6d** and **6e**). While the PacL2^Δ^^31–84^ variant appeared unstable (**Fig. 6d** and **6f**), the PacL2^Δ55–84^ variant was still able to form fluorescent foci in *M. smegmatis*.

We also generated PacL1 deletion mutants lacking segments of its cytoplasmic domain (PacL1^Δ54–86^ and PacL1^Δ37-86^) (**Fig. 6a**). In strains expressing CtpC-mVenus fusion proteins, fluorescent foci observed with wild-type PacL1 co-expression were absent when co-expressed with these PacL1 deletion mutants (**Fig. 6g** and **6h**). However, despite a reduced ability to form large clusters, both deletion variants remained stable and continued to form small fluorescent foci in the mycobacterial membrane (**Fig. 6d** and **6f**). Overall, these results suggest that the cytoplasmic domain of PacL proteins, including the AE repeats, is not the primary determinant of PacL cluster formation, although it plays a crucial role in stabilizing interactions with P-type ATPase pumps and ensuring proper protein function.

### Conserved amino acids of the TM domain of PacL proteins are essential for assembling functional multi-metal efflux machineries

To identify critical amino acids in the TM domain of PacL proteins essential for PacL/PacL interactions, we aligned 120 sequences of PacL-like proteins identified in our previous study across multiple bacterial species^5^. The most highly conserved residues included a lysine (K9) and two glycines (G17 and G20), separated by two variable amino acids (**Fig. 7a** and **S5a**). Notably, the two glycines form a groove in the TM domain (**Fig. S5b**) resembling the GXXXG motif, a well characterized sequence that facilitates helix-helix interactions in membrane proteins^18^.

**Figure 7.**
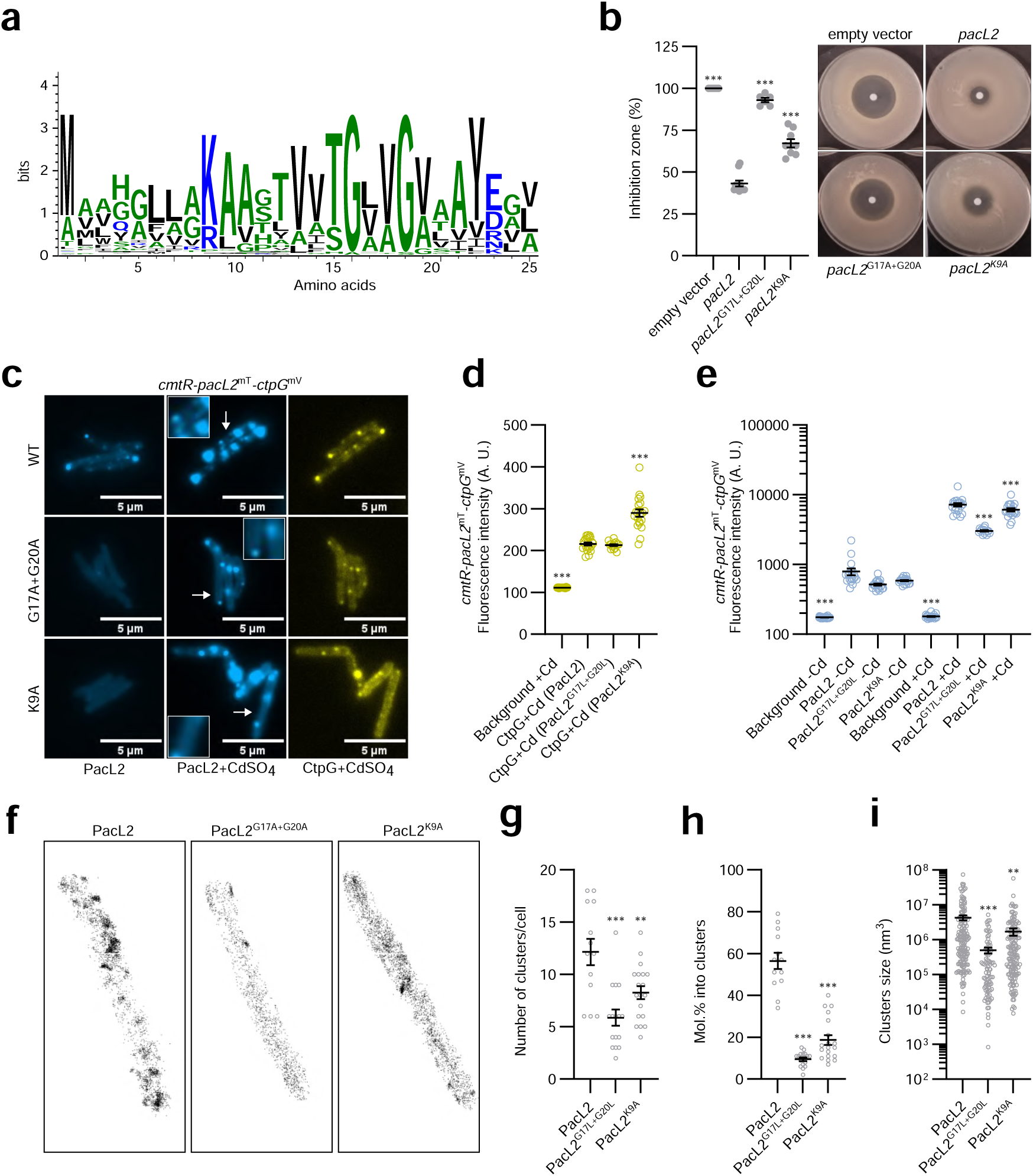
Conserved amino acids of the TM domain of PacL proteins are essential to assemble functional PacL clusters. (**a**) Sequence conservation logo of the predicted transmembrane domain of 120 bacterial PacL proteins. (**b**) Sensitivity of the indicated *M. smegmatis* strains to CdSO_4_, assessed by disk diffusion assay. Results represent the diameter of the inhibition zone, expressed relative to the empty vector condition. Data are presented as means (±SEM) from biological replicates, represented by gray dots. (**c**) Epifluorescence microscopy images of live *M. smegmatis* cells expressing the indicated fusion proteins under their respective native promoters cultivated in 7H9 medium supplemented or not with 100 nM CdSO_4_. (**d, e**) Fluorescence intensity measurements of the indicated strain in grayscale. Results are presented as means (±SEM) from individual cells, represented by yellow or blue dots. (**f**) Super-resolution 3D imaging of individual PacL2-mEos proteins (PacL2, PacL2^G17L+G20L^, or PacL2^K9A^) using PALM microscopy in *M. smegmatis* cells cultivated in 7H9 medium supplemented with 100 nM CdSO_4_. Representative images are shown from dozens of analyzed cells. (**g**–**i**) Quantification of (**g**) the number of PacL2 clusters per cell, (**h**) cluster sizes, and (**i**) the proportion of molecules localized within a cluster per cell, determined via cluster segmentation of single-molecule localizations using the PoCA software platform. Results are presented as means (±SEM) from individual cells (**g, i**) or individual clusters (**h**), represented by gray dots. Asterisks indicate statistically significant differences compared to the reference condition (empty vector (b) or WT (**d, e, g, h, i**) (**P < 0.01; ***P < 0.001).

To investigate the role of these conserved residues in PacL/PacL interactions, we constructed *M. smegmatis* strains expressing PacL2 variants with substitutions of these residues (PacL2^G17L+G20L^ and PacL2^K9A^) (**Fig. S5b**). To assess the functional impact of these mutations, we evaluated PacL2/CtpG-dependent cadmium tolerance using a disc diffusion assay. Both mutations significantly reduced cadmium tolerance (**Fig. 7b**), indicating the importance of these residues for proper function. Interestingly, epifluorescence microscopy showed no reduction in fluorescence intensity in cells expressing CtpG-mVenus co-expressed with either PacL2, PacL2^G17L+G20L^, or PacL2^K9A^ (**Fig. 7c**, and **7d**). This suggests that while these mutations disrupt CtpG functionality, they do not impair the ability of PacL2 to stabilize the P-type ATPase pump directly.

Interestingly, although the fluorescence intensity of PacL2-mTurquoise was weakly affected by the G17L+G20L and K9A mutations (**Fig. 7e**), we observed significant differences in cluster formation. In the absence of cadmium, while wild-type PacL2 formed multiple distinct fluorescent foci, the majority of cells expressing the PacL2^G17L+G20L^ and PacL2^K9A^ mutants exhibited no detectable foci (**Fig. 7c** and <u>Movie 1</u>, <u>Movie 3</u> and <u>Movie 4</u>). In the presence of cadmium, both mutants were still able to form large, immobile foci that colocalized with CtpG clusters. However, the number of small, mobile clusters, characteristic of wild-type PacL2, was strongly reduced in the PacL2^G17L+G20L^ and PacL2^K9A^ mutants (**Fig. 7c** and <u>Movie</u> <u>1</u>, <u>Movie 3</u>, and <u>Movie 4</u>). Additionally, the fluorescence signal of both mutants appeared more diffuse throughout the cells, suggesting that these mutations disrupt the ability of PacL2 to organize into distinct membrane clusters, resulting in a more homogeneous distribution of PacL proteins within the membrane.

To gain deeper insights into cluster formation in the PacL2^G17L+G20L^ and PacL2^K9A^ mutants, we constructed fusion proteins with mEos for super-resolution microscopy (**Fig. 7f**). Cluster segmentation of 3D localizations revealed that PacL2^G17L+G20L^ and PacL2^K9A^ formed an average of 6 and 8 clusters per cell, respectively, compared to 12 clusters in wild-type PacL2 (**Fig. 7f** and **7g**). Additionally, both mutant proteins were more homogeneously distributed across the membrane, with only 10% and 19% of PacL2^G17L+G20L^ and PacL2^K9A^ molecules localizing within clusters, compared to 57% for the wild-type protein (**Fig. 7f** and **7h**). Furthermore, the clusters formed by both mutants were significantly smaller than those formed by wild-type PacL2, with average volumes of 5.10[nm³ and 2.10[nm³ for the mutants, compared to 4.10[nm³ for the wild-type (**Fig. 7f** and **7h**). Altogether, these findings indicate that the GXXG motif and the K9 residue of PacL2 trans-membrane segment are essential for cluster formation. Moreover, they suggest that the function of CtpG depends on its localization within small, mobile clusters, rather than being associated with large, immobile aggregates or a homogeneously distributed membrane state.

### Interaction network of PacL1 with *M. tuberculosis* proteins

The results presented above reveal that PacL proteins act as hub molecules, organizing multiple P-ATPases into membrane-associated platforms. To investigate whether PacL-dependent clusters are limited to CtpC, CtpV, and CtpG or also include additional interacting proteins, we utilized a recently developed split ALFA tag nanobody system for proximity labeling^19^ to map the PacL1 interaction network in *M. tuberculosis*. For this analysis, we expressed a PacL1 protein fused to a C-terminal or an internal ALFA tag^20^ (PacL1^Cter-ALFA^ and PacL1^Int-ALFA^) in an *M. tuberculosis* Δ*pacL1* mutant, alongside a TurboID^21^-nanobody fusion protein designed to bind the ALFA tag and biotinylate proteins in close proximity^19^. The internal ALFA tag was incorporated between A^34^ and P^35^ of PacL1, downstream of its TM domain. Immunoblots showed that constructed *M. tuberculosis* strains properly expressed tagged PacL1 proteins and TurboID (**Fig. S6a**). Importantly, co-expression of PacL1^Cter-ALFA^ and PacL1^Int-ALFA^ with CtpC and the TurboID-nanobody fusion conferred zinc tolerance in both *M. smegmatis* (**Fig. S6b** and **c**) and *M. tuberculosis* (**Fig. S6d**), confirming that the binding of the fusion protein to PacL1 does not abolish the formation or functionality of PacL-dependent membrane clusters and that the ALFA tag does not disrupt PacL1 function.

Biotinylated proteins were identified by proteomics and compared to those detected under control conditions, where the TM domain of the *E. coli* MalF protein carrying an ALFA-tag (MalF_(1,2)-_ALFA) was co-expressed with the TurboID-nanobody fusion protein. This control protein is uniformly distributed across the mycobacterial membrane^22^, providing a baseline for non-specific interactions. We respectively identified 46 and 22 proteins that were at least twice as biotinylated in the strain expressing PacL1^Cter-ALFA^ and PacL1^Int-ALFA^ compared to the strain expressing MalF_(1,2)-_ALFA (**Table 1**). Among them, seven were enriched in both PacL1^Cter-αtag^ and PacL1^Int-ALFA^ strains: PacL1, Rv1265, CtpC, PPE20, MoeX, Rv2083, and AhpC. According to Gene Ontology (GO) annotations from QuickGO, among the 61 identified proteins, 25 are predicted to localize to the plasma membrane or cell wall, 7 to the cytoplasm, 4 to extracellular compartments, while the subcellular localization of the remaining proteins remains unknown. Notably, PacL1, CtpC, CtpV, and PacL3 were found enriched confirming that they are part of the effluxosome. The absence of PacL2 and CtpG from the identified proteins is likely due to their low expression levels, as the experiment was conducted without cadmium supplementation, which typically induces their expression.

**Table 1.** Network of PacL1 interactions with *M. tuberculosis* proteins. *M. tuberculosis* proteins significantly more biotinylated (Log2 fold-change > 1;-log10 p-value > 1.3) in a strain expressing a TurboID-nanobody fusion protein along with the PacL1 protein containing a C-terminal ALFA tag (PacL1^Cter-ALFA^) or an internal α-tag (PacL1^Int-ALFA^), compared to a strain expressing the TurboID-nanobody fusion protein with the TM domain of the *E. coli* MalF protein carrying an ALFA-tag (MalF_(1,2)-_ALFA).

| Genes | ProteinDescriptions | Fold change (log2) | p-value (-log10) |
| --- | --- | --- | --- |
| *pacL1\** | DUF1490 family protein | 6,34 | 5,91 |
| *Rv1265\** | Uncharacterized protein | 3,74 | 4,40 |
| *ctpV* | Metal cation transporter P-type ATPase | 2,29 | 3,33 |
| *ctpC\** | Metal cation transporter P-type ATPase | 2,08 | 5,12 |
| *PPE20\** | PPE family protein | 1,93 | 4,29 |
| *efp* | Elongation factor P | 1,65 | 2,34 |
| *Rv3898c* | Uncharacterized protein | 1,58 | 2,45 |
| *Rv3559c* | Short chain dehydrogenase | 1,54 | 3,28 |
| *citE* | Citrate (Pro-3S)-lyase beta subunit | 1,53 | 3,69 |
| *Rv2302* | DUF1918 domain-containing protein | 1,52 | 1,42 |
| *Rv2114* | Uncharacterized protein | 1,47 | 2,26 |
| *Rv0123* | DNA-binding protein | 1,44 | 1,50 |
| *pacL3* | DUF1490 family protein | 1,44 | 4,58 |
| *Rv3357* | Antitoxin | 1,44 | 3,91 |
| *Rv3878* | ESX-1 secretion-associated protein EspJ | 1,43 | 2,65 |
| *Rv1637c* | Metallo-beta-lactamase superfamily protein | 1,37 | 2,15 |
| *moeX\** | Molybdopterin biosynthesis protein | 1,36 | 2,45 |
| *sahH* | Adenosylhomocysteinase | 1,34 | 3,47 |
| *Rv2857c* | Short chain dehydrogenase | 1,33 | 3,99 |
| *hns* | Histone-like protein | 1,32 | 2,66 |
| *TB9.4* | ATP-binding protein | 1,30 | 1,83 |
| *Rv3788* | Nucleoside diphosphate kinase regulator | 1,26 | 1,85 |
| *rpmG* | Large ribosomal subunit protein bL33 | 1,24 | 2,12 |
| *Rv2083\** | PPE family protein | 1,22 | 1,53 |
| *greA* | Transcription elongation factor GreA | 1,21 | 2,21 |
| *Rv0121c* | Pyridoxamine 5'-phosphate oxidase | 1,21 | 2,46 |
| *PPE41* | Uncharacterized protein | 1,20 | 2,90 |
| *Rv0250c* | Uncharacterized protein | 1,19 | 4,26 |
| *mog* | Molybdopterin biosynthesis Mog protein | 1,19 | 3,63 |
| *Rv2830c* | Uncharacterized protein | 1,16 | 2,02 |
| *Rv1566c* | Inv protein | 1,16 | 2,39 |
| *Rv3474* | Probable transposase | 1,15 | 3,93 |
| *ahpC\** | Alkyl hydroperoxide reductase subunit C | 1,15 | 4,84 |
| *cspA* | Cold shock protein A | 1,12 | 3,25 |
| *Rv3697A* | CopG family DNA-binding protein | 1,10 | 2,14 |
| *argR* | Arginine repressor | 1,10 | 2,45 |
| *esxB* | ESAT-6-like protein | 1,08 | 2,27 |
| *Rv3881c* | Uncharacterized protein | 1,07 | 3,14 |
| *galU* | UTP--glucose-1-phosphate uridylyltransferase | 1,07 | 3,11 |
| *fadD30* | Acyl-CoA synthetase | 1,04 | 2,85 |
| *hbhA* | Iron-regulated heparin binding hemagglutinin | 1,03 | 3,56 |
| *Rv3421c* | Gcp-like domain-containing protein | 1,01 | 2,07 |
| *Rv3046c* | DUF3349 domain-containing protein | 1,01 | 2,59 |
| *frdA* | Fumarate reductase flavoprotein subunit | 1,01 | 1,83 |
| *Rv0887c* | VOC domain-containing protein | 1,01 | 2,44 |
| *mihF* | Integration host factor | 1,01 | 1,30 |

\*proteins detected in both conditions

**Table 1. Network of PacL1 interactions with *M.**
| Genes | ProteinDescriptions | Fold change (log2) | p-value (-log10) |
| --- | --- | --- | --- |
| *pacL1\** | DUF1490 family protein | 6,91 | 8,26 |
| *Rv1265\** | Uncharacterized protein | 4,48 | 6,06 |
| *ctpC\** | Metal cation transporter P-type ATPase | 3,20 | 7,33 |
| *Rv2304c* | Uncharacterized protein | 2,47 | 4,83 |
| *PPE20\** | PPE family protein | 2,44 | 4,08 |
| *Rv1066* | Rhodanese domain-containing protein | 1,69 | 1,63 |
| *Rv2083\** | PPE family protein | 1,61 | 2,91 |
| *Rv2295* | CbrC family protein | 1,51 | 4,49 |
| *Rv0572c* | Uncharacterized protein | 1,51 | 2,38 |
| *Rv2204c* | FeS cluster biogenesis domain-containing prot. | 1,50 | 4,35 |
| *fdxA* | Ferredoxin | 1,33 | 1,75 |
| *Rv0680c* | Transmembrane protein | 1,29 | 3,07 |
| *Rv0784* | Deacetylase | 1,18 | 2,38 |
| *ruvC* | Crossover junction endodeoxyribonuclease | 1,17 | 2,90 |
| *moeX\** | Molybdopterin biosynthesis protein | 1,11 | 1,53 |
| *Rv2809* | Uncharacterized protein | 1,07 | 2,61 |
| *Rv3877* | Transmembrane protein | 1,05 | 3,19 |
| *Rv2686c* | Antibiotic ABC transporter | 1,04 | 2,58 |
| *Rv0133* | Acetyltransferase | 1,03 | 5,30 |
| *chdC* | Coproheme decarboxylase | 1,02 | 4,52 |
| *rpe* | Ribulose-phosphate 3-epimerase | 1,01 | 2,01 |
| *ahpC\** | Alkyl hydroperoxide reductase subunit C | 1,01 | 4,32 |

The analysis revealed a diverse array of proteins with various functions. Among them, we identified several PE/PPE family proteins, including PPE20, PPE41, and Rv2083, as well as components of the ESX-1 secretion system, such as EsxB (CFP-10), Rv3877, Rv3878, and Rv3881c, which are known to play roles in *M. tuberculosis* virulence and immune modulation^23,24^. Additionally, several proteins, including AhpC, CspA, and GreA, are implicated in *M. tuberculosis* adaptation to environmental stress^25–27^. Finally, the identification of multiple proteins with unknown functions, such as Rv1265 which was strongly enriched in both PacL1^Cter-ALFA^ and PacL1^Int-ALFA^ strains, suggests the presence of potential novel players in metal detoxification and/or stress adaptation. These findings indicate that the effluxosome could extend beyond CtpC, CtpG, and CtpV, encompassing a broader network of proteins that may enhance *M. tuberculosis*’s ability to withstand environmental stresses and contribute to its pathogenic potential.

## Discussion

In this study, we found that CtpC and CtpG confer cross-resistance to zinc and cadmium, while the function of CtpV remains unclear. We showed that PacL1, PacL2, and PacL3 cluster in the mycobacterial membrane *via* transmembrane CXXC motifs and recruit CtpC, CtpG, and CtpV through EA repeats, forming dynamic membrane complexes essential for P-ATPase function. We demonstrated that PacL1 binds multiple metals, enhancing P-ATPase-mediated tolerance. For the first time, we reveal that multiple P-ATPase pumps assemble into dynamic membrane-associated platforms, which we term effluxosomes (**Fig. 8**), raising important questions about the biological significance of these structures.

**Figure 8.**
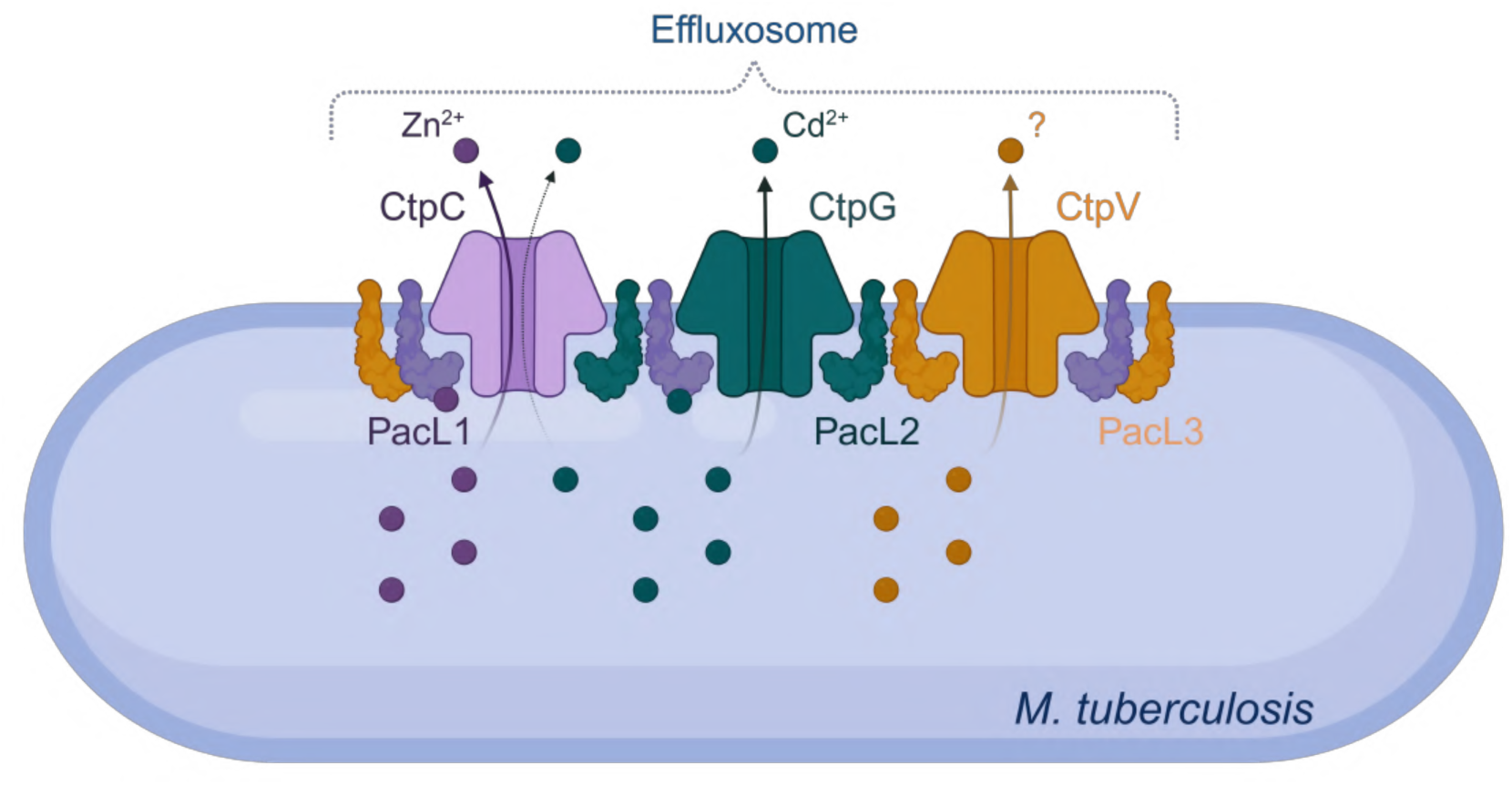
Conceptual model of mycobacterial effluxosome for multi-metal cross-resistance. The P-ATPases CtpC and CtpG facilitate the efflux of zinc and cadmium from the cell, while the role of CtpV remains unclear. The PacL1, PacL2, and PacL3 proteins interact with each other through the CXXC motif in their transmembrane domains and with the metal-binding domains of CtpC, CtpG, and CtpV via EA repeat sequences in their cytoplasmic domains. These interactions are essential for the activity of the P-ATPases, clustering them into dynamic mobile membrane complexes. PacL1, but not PacL2, binds a variety of metal ions, thereby enhancing the metal tolerance conferred by the P-ATPases.

Building on previous research, we recently discussed various examples of bacterial membrane protein that form clusters, highlighting the potential advantages of such assemblies within membranes^28^. These benefits include promoting oligomerization, protecting proteins from degradation, and facilitating preferential subcellular organization. Here, our findings indicate that the clustering of P-ATPase pumps by PacL proteins not only shields them from degradation but is also essential for their functionality, although further studies are needed to elucidate the underlying mechanisms. Another intriguing hypothesis regarding the biological relevance of clustering different P-ATPases relates to the ability of PacL1 to bind multiple metals. This activity could locally concentrate various metal ions, potentially enhancing the efflux efficiency of P-ATPases specialized in exporting specific ions. However, whether direct metal transfer occurs between PacL1 and individual P-ATPases remains to be determined.

To explore the broader relevance of these findings, we examined the distribution of PacL proteins across bacterial species. In a previous study, we identified 98 bacterial species encoding at least one PacL protein (containing a DUF1490 domain), all of which belong to the Actinobacteria phylum. The majority of these species (84%) harbor a single *pacL* gene, which is constantly in operon with a gene encoding a P-ATPase pump. Interestingly, 10% of these species encode two PacL/Ctp pairs, 4% encode three pairs, and 2% encode five pairs. This distribution suggests that effluxosomes may not be unique to *M. tuberculosis* but could also be present in other pathogenic or opportunistic bacteria, such as *Mycobacterium marinum*, which notably encodes five PacL/Ctp pairs. PacL proteins share close similarity with DUF6110-containing proteins, which we refer to as PacL-like proteins. These homologues are primarily found in Firmicutes but also occur in other bacterial phyla, including Actinobacteria and Proteobacteria, as well as in Archaea. With few exceptions, these organisms encode only a single PacL-like protein. However, since our study demonstrates that each PacL protein can interact with multiple P-ATPase pumps, we cannot exclude the possibility that a single PacL protein might associate with P-ATPases encoded elsewhere in the genome. This raises the intriguing possibility that effluxosome-like assemblies could form even in bacteria encoding only one PacL or PacL-like protein.

Given the role of P-ATPases in metal tolerance, we next investigated the specific contribution of CtpG. Previous studies have reported that CtpG contributes to tolerance against zinc, cadmium, and copper^9,14^. In our study, we assessed the ability of CtpG to confer tolerance to multiple metals, including cadmium, zinc, copper, nickel, manganese, and iron. Our findings reveal that CtpG specifically plays a role in cadmium tolerance, but does not contribute to resistance against the other metals. This discrepancy with previous reports could, in part, be explained by the requirement of PacL2 for CtpG function, as enzymatic activity measured in isolation from the P-ATPase may not reflect optimal physiological conditions.

The relevance of cadmium tolerance in *M. tuberculosis* infection remains an open question. Although cadmium has no known biological function in bacteria, it interacts with nucleic acids, competes with essential metal cofactor in proteins, and induces oxidative stress by promoting reactive oxygen species production^29^. However, its potential role in pathogen clearance by the immune system remains unexplored. Notably, *ctpG* expression is induced during macrophage infection^15^ and has been shown to contribute to bacterial growth within the host^9^. The cadmium specificity of CtpG raises the intriguing question of whether *M. tuberculosis* encounters toxic cadmium concentrations during infection. Cadmium, largely derived from environmental pollution and cigarette smoke, accumulates in alveolar macrophages^30^, the natural niche of *M. tuberculosis*, suggesting that the pathogen is likely exposed to this metal in the host. Future studies should investigate whether macrophages locally elevate cadmium concentrations in the phagosome to intoxicate *M. tuberculosis*, as has been previously reported for zinc^2^.

Beyond cadmium, we also examined the role of CtpV, a P-ATPase previously implicated in copper tolerance. A previous study demonstrated that an *M. tuberculosis ctpV* mutant exhibits increased sensitivity to copper, suggesting that CtpV plays a role in copper efflux^10^. This would be consistent with the fact that CtpV clusters with copper efflux pumps in phylogenetic analyses^3^. However, in this study, we did not observe an impact of *ctpV* expression in *M. smegmatis* on tolerance to copper or other metals. This finding is particularly unexpected given the strong induction of the *ctpV* operon in response to copper^10,31^. Although we did not detect increased copper sensitivity in a *M. tuberculosis* triple *pacL* mutant (data not shown), we cannot rule out the possibility that CtpV requires an additional, yet unidentified, essential partner encoded in the *M. tuberculosis* genome for its function. Alternatively, CtpV may not primarily function in copper detoxification but instead facilitate copper transport to deliver it as a cofactor to periplasmic and membrane-associated proteins, as has been proposed for CtpA^32^ and CtpB^33^. Future studies will be needed to clarify the precise role of CtpV in *M. tuberculosis* and its potential involvement in effluxosome function.

Interestingly, our findings suggest that the role of the effluxosome extends beyond metal detoxification. Several proteins interacting with PacL1 appear to have broader functions in stress adaptation, immune response modulation, and virulence. Among them, Rv1265 is particularly intriguing, as it exhibited a 13-and a 22-fold increase in biotinylation in strains expressing PacL1^Cter-αtag^ and PacL1^Int-αtag^, respectively, compared to the control. Rv1265 is known to be induced early during macrophage infection and responds to starvation and elevated cAMP levels under hypoxic conditions^34^, suggesting a role in mycobacterial stress adaptation. Notably, Rv1265 protein has been reported to associate with the bacterial cell envelope, where it modulates cell envelope composition and enhances mycobacterial survival within macrophages^34^. However, a more recent study suggests that Rv1265 functions as an ATP-binding transcription factor regulating the expression of the small non-coding RNA Mcr11^35^. Another protein of interest is PPE20, which was enriched 4-and 5-fold in PacL1^Cter-αtag^ and PacL1^Int-αtag^ strains, respectively. Interestingly, PPE20 was recently identified as a transporter involved in calcium translocation across the *M. tuberculosis* outer membrane^36^. Future studies will be crucial to determine whether Rv1265, PPE20, and other identified proteins form stable, membrane-associated clusters in mycobacteria and to elucidate their specific functional roles within these newly identified membrane machineries.

Overall, our study provides new insights into how bacterial pathogens organize specialized membrane platforms to resist metal ion stress. The effluxosome concept refers to a unique cross-resistance strategy that enables *M. tuberculosis* to withstand stress induced by multiple metals, and possibly other compounds, encountered within the phagosome^4^. Given that CtpC, CtpG, and CtpV contribute to mycobacterial growth within the host^2,6–10^, and that other proteins, including stress response factors, may also be part of the effluxosome, our findings have important implications for the development of novel antimicrobial strategies. Our future research will investigate whether the effluxosome plays a broader role in stress adaptation beyond metal detoxification, and whether it contributes to *M. tuberculosis* virulence.

## Methods

### Bacterial Strains

The bacterial strains used in this study are listed in Supplementary Table 1. *Escherichia coli* strains were cultivated in Luria-Bertani (LB) medium at 37 °C. *Mycobacterium smegmatis* mc²155 strains (ATCC 700084) were grown in Middlebrook 7H9 medium (Difco) at 37 °C, supplemented with 0.5% glycerol, 0.5% dextrose, and 0.05% Tween 80. *M. tuberculosis* H37Rv (ATCC 27294) strains were grown at 37 °C in 7H9 medium (Difco) supplemented with 10% albumin-dextrose-catalase (ADC, Difco) and 0.05% Tween-80 (Sigma-Aldrich), or on complete 7H11 solid medium (Difco) supplemented with 10% oleic acid-albumin-dextrose-catalase (OADC, Difco). When required, streptomycin (5 μg.ml^−1^), hygromycin (50 μg.ml^−1^) or zeocin (25 μg.ml^−1^) were added to the culture media.

### Plasmids

Plasmids and oligonucleotides used in this study are listed in Supplementary Tables 2 and 3. Plasmids were constructed in *Escherichia coli* Stellar™ (Takara Bio), and cloning was performed using the In-Fusion recombination cloning kit (Takara), as recommended by the manufacturer. DNA fragments were amplified via PCR using PrimeSTAR GXL DNA polymerase (Takara Bio), plasmidic or genomic DNA from *Mycobacterium tuberculosis* H37Rv as template, and specific primer pairs listed in Supplementary Table 3. Plasmids for the expression of the *pacL* and *ctp* genes under their native promoters were generated by amplifying the open reading frames (ORFs) along with their 5′ flanking regions (∼500 bp) via PCR and inserting them into EcoRI-digested pDB60. For the ATC-inducible plasmids, ORFs were amplified without 5′ flanking regions and cloned into ClaI-digested pMSG419. Plasmids for bi-partite split GFP were constructed by cloning the ORFs of the proteins of interest (PacL, MBD of Ctp, or Rv1488) along with the GFP N-terminal fragment (GFP1-10) or the GFP C-terminal fragment (GFP11), separated by a linker sequence, into the pGMC vector. Derivative plasmids containing deletions, substitutions, or tagged versions were constructed by In-Fusion cloning using the appropriate primer pairs (Supplementary Table 3). The absence of mutations in the constructs was verified by DNA sequencing.

### Generation of *M. tuberculosis* mutants

Mutant strains of *M. tuberculosis* H37Rv were generated via allelic exchange using recombineering, following established protocols^5,37,38^. Approximately 0.5 kb DNA fragments flanking the target genes were amplified using PrimeSTAR GXL DNA polymerase (Takara Bio), genomic DNA from *M. tuberculosis* H37Rv, and specific primers (Supplementary Table 3). These fragments were fused with a zeocin-resistance cassette via a three-fragment PCR strategy, incorporating *M. tuberculosis dif* site variants for unmarked deletions. For recombineering, recipient strains of *M. tuberculosis* H37Rv, harboring pJV53H encoding recombineering enzymes^37^, were cultured to mid-log phase in 7H9 medium. Recombineering enzymes were induced with 0.2% acetamide, followed by electrotransformation with 100 ng of linear AES for allelic exchange. After 48 hours at 37°C, zeocin-resistant clones were selected and expanded. PCR analysis confirmed successful allele replacement. The *dif* site-flanked zeocin-resistance cassette was excised via spontaneous XerCD-mediated recombination, and the pJV53H plasmid was lost by serial passaging in antibiotic-free medium, followed by phenotypic screening for sensitivity to antibiotics, as described previously^38^.

### Disc Diffusion Assay

Bacteria were grown to exponential phase, diluted to OD = 0.01 in 3 ml of prewarmed top agar (7H9 with 6 mg/ml agar), and plated on 7H10 agar. A filter paper disc was placed on the solidified top agar and spotted with 2.5 μl of 1M ZnSo_4_, CdSo_4_, CuSo_4_, NiSo_4_, MnCl_2_, FeSo_4_, or CuCl_2_. After incubation at 30 °C for 72 hours, the diameter of the growth inhibition zone was measured.

### *M. tuberculosis* cadmium sensitivity Assay

*M. tuberculosis* cultures were initiated at an OD of 0.01 in glass tubes containing 7H9 medium supplemented with varying concentrations of CdSO. The cultures were incubated statically at 37 °C, and bacterial growth was assessed over time by measuring turbidity in McFarland units using a Densimat device (BioMérieux).

### Bipartite split GFP

Log phase cultures of *M. smegmatis* strains transformed with relevant plasmids were grown in 7H9 medium containing 5 µg.ml^-1^ streptomycin to an OD of 0.05. After overnight incubation at 37 °C, the bacterial cells were analyzed using fluorescence-activated cell sorting (FACS). FACS was performed using an LSR Fortessa (BD Biosciences) flow cytometer. The GFP fluorescence signal was processed with FlowJo (v10) software. Gating strategy is displayed in Supplementary Fig. S7.

### Fluorescence microscopy

Fluorescence microscopy was performed on mycobacterial cultures harboring plasmids expressing fluorescent reporters fused to the proteins of interest cultivated in the absence or presence of various metals. When necessary, mycobacterial cells were fixed with 4% paraformaldehyde (PFA) treatment (2 hours for *M. tuberculosis* and 20 minutes for *M. smegmatis*), followed by three PBS washing steps. Cultures were deposited as 1 μL drops onto a 1% agarose layer in growth medium, following previously described method^39^. Imaging was conducted using an Eclipse TI-E/B wide-field epifluorescence microscope equipped with a phase contrast objective (CFI Plan APO LBDA 100X oil NA 1.45) and Semrock filters for YFP (Ex: 500BP24; DM: 520; Em: 542BP27), CFP (Ex: 438BP24; DM: 458; Em: 483BP32), or FITC (Ex: 482BP35; DM: 506; Em: 536BP40). Images were captured with an Andor Neo SCC-02124 camera, using 30-50% illumination from a SpectraX LED light source (Lumencor) and 0.3-0.6 second exposure time depending on the fluorochrome used. Time-lapses were performed over a 45-second period, with an image captured every second. Image acquisition and analysis were performed using Nis[Elements AR software (Nikon) and ImageJ (v 1.53f51). Fluorescence intensity measurements were obtained from individual cells using the grayscale mean intensity of the ROI, as measured with the ImageJ analysis tool.

### Photoactivated localization microscopy (PALM) and clusters analysis

PALM was performed using an inverted Nikon Ti-E/B microscope equipped with the “Perfect Focus System” (PFS, Nikon), a piezo stage (Nano Z100-N - Mad City Labs), a CFI Apochromat TIRF 100X Oil Objective (NA 1.49), and a quad-band dichroic mirror (97335 Nikon N-STORM TIRF Filter Set). A TIRF illumination arm was used, specifically in a highly inclined and laminated optical sheet (HILO) illumination mode. The system included an EM-CCD ANDOR iXon Ultra DU897 camera and a thermostatic chamber set at 24°C to minimize instrumental drift. Excitation was controlled with an AOTF. For mEos-EM photoconversion, we used a 405 nm laser (Coherent™ - Cube 405-100C), while imaging was performed with a 561 nm laser (Coherent™, Sapphire) at a power of approximately 0.1 kW/cm². The 405 nm laser was applied at low power to enhance the photoconversion rate. The imaging laser (561 nm) was used continuously, with an acquisition time of 30 ms per frame (between 10,000 and 20,000 images). Images were acquired using Nis-Elements AR software (Nikon). Point-spread function (PSF) astigmatism was introduced using adaptive optics with a deformable mirror (MicAO 3DSR – Imagine Optic™) placed in the detection path, just before the EM-CCD camera. This setup enabled axial localization of single fluorescent molecules. PSF calibrations were performed using commercial fluorescent beads (FluoSpheres™ Carboxylate-Modified Microspheres, 0.17 µm, orange fluorescent (540/560), 2% solids, Invitrogen™).

3D localization was performed using ThunderSTORM, an ImageJ plugin^40^. For each frame in the image sequence, raw images were filtered using a wavelet filter (B-Spline, order 3, scale 2.0). Approximate molecular detections were identified using the local maximum method, followed by sub-pixel localization using the PSF elliptical Gaussian method (Fitting Radius: 6 pixels, Initial Sigma: 1.6). The plugin allowed data filtering based on specific criteria. In this study, we eliminated localizations from the first 1,000 frames and retained only molecules detected with a localization precision of less than 50 nm. Strict environmental controls were applied to minimize sample drift, including air conditioning, a thermostatic chamber, the Nikon Perfect Focus System, and an x/y-controlled stage. Residual drift was estimated using an autocorrelation algorithm in ThunderSTORM software^40^. SMLM dataset visualization was also performed using ThunderSTORM. The voxel size was determined to ensure three pixels per resolution unit, and localizations were blurred using a theoretical PSF corresponding to the expected resolution.

To analyze the cluster organization of our protein of interest, we used the 3D DBSCAN module of PoCA (Point Cloud Analyst) software^17^. PoCA utilizes 3D localizations coordinates computed from 3D PALM microscopy to compute clusters based on the localization density within a given sphere (minimum localizations: 2; minimum number of localizations per cluster: 0.5% of total localizations; maximum distance between two localizations: 40 nm).

sptPALM acquisitions were performed on the same microscopy setup as the PALM acquisitions described above, using the 561 nm laser at a reduced power of approximately 0.1 kW/cm² to visualize the trajectories. The imaging laser (561 nm) was used continuously, with an acquisition time of 30 ms per frame during 20,000 images. Single molecule localization and analysis was performed using the PALM_Tracer module^41^, that use a combination of wavelet filtering and Gaussian fitting for the localization^42^, and a simulated annealing for the tracking. We used a threshold of 18 and a maximum reconnection distance of 4 pixels, corresponding to a maximum instant speed of 20 µm.s^-1^, in order to maximize the chances to capture the fast moving molecules. Reconnection artefacts were minimized thanks to the very low activation level of mEOS molecules. Diffusion coefficients (D) were computed by fitting the first fours points of the mean square displacement (MSD) function^43^.

### Expression and purification of SolPacL1 and SolPacL2

The soluble domain of PacL1 was expressed and purified as reported previously^5^. The gene of the soluble domain of PacL2 (SolPacL2) was codon optimized and cloned in pExp-GST. After amplification and purification with QIAprep Spin Miniprep Kit, the plasmid was subsequently transformed into BL21 Star (DE3). The production of unlabeled SolPacL2 was performed in Luria Broth medium supplemented with 100 µg/mL Amp in 1L flask at 37 °C under vigorous shaking at 170 rpm. SolPacL2 expression was induced during 4 hours by the addition of 1 mM isopropyl-β-D-thiogalactopyranoside (IPTG) at 37 °C. Then, cells were harvested by centrifugation (5,000 x g, 10 min, 4 °C) and stored at-70 °C. The cells were suspended in 5 to 10 mL of lysis buffer per gram of wet cell (50 mM Tris-HCl pH 7.5, 200 mM NaCl, 0.5 % Tween20, 1 mM phenylmethanesulfonyl fluoride (PMSF), 1 mM Dithiothreitol (DTT), DNAse (0.01-1mg/mL) and cOmplet^TM^ EDTA-free protease inhibitor cocktail. Cells were lysed by passing the suspension through the Avestin® Emulsiflex-C5 (3 times at 15,000 psi) at 4°C. Lysate was centrifuged (12,000 x g, 30 min, 4 °C). The supernatant was filtered on a 0.22 µm membrane filter and loaded at 1 mL/min onto 2 x 5 mL GSTrap^TM^ FF column (Cytiva) conditioned in the equilibration buffer (50 mM Tris-HCl pH 7.5, 200 mM NaCl, 1 mM DTT). The column was washed with 10 CV of equilibration buffer supplemented with 0.5 mM PMSF and cOmplet^TM^ EDTA-free protease inhibitor cocktail. The protein was eluted with 4 to 5 CV of elution buffer (50 mM Tris pH 8.0, 10 mM reduced glutathione). The 8-His-GST tag was cleaved by addition of TEV protease at a mass ratio 1:50 in the elution buffer supplemented with 500 µM EDTA, 1 mM DTT, 100 µM PMSF. The cleavage reaction was performed overnight at 4 °C in a Spectra/Por®3 dialysis membrane (MWCO: 3.5 kDa) against 2 L of dialysis buffer (50 mM Tris-HCl pH 8.0, 125 mM NaCl, 100 µM PMSF, 500 µM DTT). The cleaved SolPacL2 was concentrated using a 3,000 MWCO PES Vivaspin (Cytiva) at 4°C, then loaded on Superdex 75 Increase 10/300 GL column (Cytiva) and eluted (V_E_∼12.5 mL) with a buffer containing 25 mM MES pH 7.0, 200 mM NaCl. At each step of the purification process the sample was characterized on SDS-Page Gel (5-10%). The concentration of the purified SolPacL2 were determined by integration (e.g. 1.0 to 0.82 ppm) of the ^1^H methyl groups of I,L,V residues by ^1^H NMR with respect to the 116.6 µM of sodium salt of 3-(Trimethylsilyl)-1-propanesulfonic acid sodium salt (DSS) used as internal reference. For NMR purpose, both ^15^N-labelled SolPacL1 and SolPacL2 expression and purification were performed in similar conditions excepted that cells were grown of minimal M9 medium supplemented with 100 µg/mL Amp, 5 g/L of glucose, 2 g/L of NH_4_Cl ^15^N, 2 mM MgSO_4_, 0.1 mM CaCl_2_, trace elements and vitamins^44^.

### Titration of Zinc and Cadmium by NMR spectroscopy

NMR experiments were performed on 600 Avance III HD spectrometer (Bruker) equipped with a 5 mm triple resonance cryoprobe at 280 K in 25 mM MES buffer pH 7.0, 200 mM NaCl and 126.6 µM DSS as chemical shift reference. The solution of ^15^N-labelled PacL1 at 90 µM was titrated with 0, 0.2, 0.4, 0.65, 1, 1.5 and 2 equivalents of ZnCl_2_ and CdCl_2_. ^1^H,^15^N-HSQC NMR data were acquired with 1024 and 256 data points respectively for ^1^H and ^15^N dimensions (e.g. 70.7 ms and 95.6 ms acquisition times respectively) and with 8 scans. All residues were assigned previously at 700 MHz (Avance III HD Bruker) on a 124 µM ^15^N-^13^C-labelled SolPAcL1 in 10 mM MES buffer pH 6.5 and 100 mM NaCl. The sequential backbone resonance assignment was performed using the best version of HNCACB, HN(CO)CACB, and HNCO experiments^45^ with selective ^1^H pulses centered at 8.5 ppm, covering a bandwidth of 4.0 ppm, and a standard HN(CA)CO experiment. NMR spectra were processed using Topspin 4.1.4 and resonances assignment was performed with cara^46^. The chemical shift perturbations (CSP) on ^15^N-labelled PacL1 induced by the interaction of zinc and cadmium with ^15^N-labelled PacL1 were classically determined with a=0.1^47^ (or B=9.9) using Sparky 3.190 (<u>Sparky - NMR Assignment Program</u>).

### Native mass spectrometry

Prior to native MS analysis, SolPacL samples at 100 µM was desalted in 200 mM ammonium acetate, pH 7, supplemented with 500 µM of zinc acetate dihydrate, manganese acetate tetrahydrate, copper (II) acetate hydrate or cadmium iodide (all from Euromedex, Souffelweyersheim, France) using Micro Bio-Spin devices (Bio-Rad, Marnes-la-Coquette, France). Samples were analyzed on a Synapt G2-Si mass spectrometer (Waters Scientific, Wilmslow, UK) running in sensitivity mode, positive ion mode and coupled to an automated chip-based nano-electrospray source (Triversa Nanomate, Advion Biosciences, Ithaca, NY). The voltage applied to the chip and the cone voltage were set to 1.6 kV and 20 V, respectively. The instrument was calibrated with a 2-mg/mL cesium iodide solution in 50% isopropanol. Raw data were acquired in the 1,000–8,000 m/z range with MassLynx 4.1 (Waters, Manchester, UK) and deconvoluted with UniDec 4.4.0^48^ using the following parameters: m/z range: 500–2,000 Th; Gaussian smoothing: 10; charge range: 1–15; mass range: 5,000–10,000 Da; sample mass every 1 Da; smooth charge states distributions; smooth nearby points: some; suppress artifacts: none; peak detection range: 22 Da, and peak detection threshold: 0.05.

### Proximity labeling

Strains for proximity labeling were prepared as follows. Cells were pre-grown in 7H9 OADC supplemented with 20 mg/ml kanamycin and 20 mg/ml streptomycin to an OD600 of 1.5-2.0 in roller bottles (Corning, # 430195). Cells were then pelleted by centrifugation at 3700xG for 10in, washed once with 10mL of biotin free Sauton’s media, subculture into biotin free Sauron’s media maintaining kanamycin and streptomycin selection, 3×25ml cultures/strain, in 60ml inkwell bottles (Nalgene, #342020-060) at a final OD600 of 0.4. ATc (50ng/ml) was added, and cultures were incubated at 37C for 18hrs. Following ATc induction cells were again collected by centrifugation at 3700xG for 10min at room temperature, resuspended in 2ml antibiotic and ATc free Sauton’s media, transferred to 2ml O-ring sealed tube, collected by centrifugation at 3700xG for 5min at room temperature, and finally resuspended in 1ml Sauton’s media. Cultures were incubated for 3hrs with 200mM Biotin (Sigma, B4501) with shaking at 37C. Cells were collected by centrifugation at 20,000xG for 1min at 4C and placed immediately on ice. Cell pellets were washed with 2ml cold TBS (20mM Tris, 150mM NaCl, pH 7.5), then collected by centrifugation at 20,000xG for 1min at 4C and cell pellets frozen at-80 degrees C.

For lysate preparation, frozen cell pellets were thawed on ice and resuspended with 250ul bead mix (1:1 mix, Biospec 11079101z and 1079107zx) and 1ml cold TBS with protease inhibitors (PI) (Pierce, PIA32955). Lysis was performed by mechanical disruption 3X45 seconds (Biospec Minibeadbeater) with 5 min icing between runs. Unbroken cells/debris were collected by centrifugation at 20,000xG for 1min at 4C, supernatants collected, and residual pellets lysed again with beads and 400ul cold TBS with protease inhibitors. Pooled supernatants were mixed with 150ul of 10x RIPA buffer detergents (TBS+ 10% triton x-100, 1% SDS, 5% sodium deoxycholate)[and mixed by inversion, centrifuged at 20,000xG for and (for Mtb) the supernatant was filtered twice through EMD Millipore™ Ultrafree™-CL Centrifugal Filter Devices with Durapore™ Membrane to remove any residual *M. tuberculosis*. For biotinylated protein capture, protein lysates were mixed with 80ul of Streptavidin-Agarose (GoldBio, S-105-10) prewashed 3X in 1ml TBS and incubated rotating at 4C for 18h. Agarose beads were collected at 250xG for 1min at 4C and washed 2xs with 1ml RIPA/SDS (TBS+ 0.1% triton x-100, 0.1% SDS, 0.5% sodium deoxycholate) followed by 4 washes with TBS.

For quantitation of biotinylated proteins, the beads were resuspended in 80 mL of 2M Urea, 50 mM ammonium bicarbonate (ABC) and treated with DL-dithiothreitol (DTT) (final concentration 1 mM) for 30 minutes at 37°C with shaking at 1100 rpm on a Thermomixer (Thermo Fisher). Free cysteine residues were alkylated with 2-iodoacetamide (IAA) (final concentration 3.67 mM) for 45 minutes at 25°C with shaking at 1100 rpm in the dark. The reaction was quenched using DTT (final concentration 3.67 mM). LysC (750 ng) was added, and the samples were incubated for 1h at 37°C with shaking at 1150 rpm. Trypsin (750 ng) was then added, and the digestion mixture was incubated for 16 hours at 37°C with shaking at 1150 rpm. Following this, an additional 500 ng of trypsin was added, and the samples were incubated for another 2-hours at 37°C with shaking at 1150 rpm. The digest was then acidified to pH <3 with 50% trifluoroacetic acid (TFA). Peptides were desalted using C18 stage tips (Empore C18 extraction disks). The stage tips were conditioned with sequential additions of: i) 100 mL methanol), ii) 100 mL 70% acetonitrile (ACN)/0.1% TFA, iii) 100 mL 0.1% TFA twice. After conditioning, the acidified peptide digest was loaded onto the stage tip, followed by two washes with 100 mL 0.1% formic acid (FA). Peptides were eluted with 50 mL 70% ACN/0.1% FA twice. Eluted peptides were dried under vacuum in a SpeedVac centrifuge, reconstituted in 12 μL of 0.1% FA, sonicated and transferred to an autosampler vial. Peptide yield was quantified using a NanoDrop (Thermo Fisher).

For mass spectrometry (MS) analyses, peptides were separated on a 25 cm column with a 75 mm diameter and 1.7 mm particle size, composed of C18 stationary phase (IonOpticks Aurora 3 1801220) using a gradient from 2% to 35% Buffer B over 90 minutes, followed by an increase to 95% Buffer B for 7 minutes (Buffer A: 0.1% FA in HPLC-grade water; Buffer B: 99.9% ACN, 0.1% FA) with a flow rate of 300 nL/min on a NanoElute2 system (Bruker). MS data were acquired on a TimsTOF HT (Bruker) with a Captive Spray source (Bruker) using a data-independent acquisition PASEF method (dia-PASEF). The mass range was set from 100 to 1700 m/z, and the ion mobility range from 0.60 V.s/cm^2^ (collision energy 20 eV) to 1.6 V.s/cm^2^ (collision energy 59 eV), a ramp time of 100 ms, and an accumulation time of 100 ms. The dia-PASEF settings included a mass range of 400.0 to 1201.0 Da, mobility range 0.60-1.60, and an estimated cycle time of 1.80 seconds. The dia-PASEF windows were set with a mass width of 26.00 Da, a mass overlap 1.00 Da, and 32 mass steps per cycle.

For DIA Data Analysis, Raw data files were processed using Spectronaut version 19.1 (Biognosys) and searched with the PULSAR search engine against the Mycobacterium smegmatis MC2155 database (12,661 entries downloaded on 2024/11/21). Cysteine carbamidomethylation was set as fixed modifications, while methionine oxidation, protein N-terminal acetylation, and asparagine/glutamine deamidation were specified as variable modifications. A maximum of two trypsin missed cleavages was allowed. A reversed sequence decoy strategy was used to control the peptide false discovery rate (FDR), with a 1% FDR threshold applied for identification. Differential analysis was performed using an unpaired t-test to calculate p-values, and volcano plots were generated based on log2 fold change (log2FC) and q-value (multiple testing corrected p-value using Benjamini-Hochberg method). A q-value of ≤0.05 was considered the statistically significant cut-off.

## Statistical analysis

One-way analysis of variance (ANOVA) and a Dunnett post-test were performed using Prism9 software (GraphPad) for all statistical analyses of this work. For bipartite split GFP experiments, the statistical analysis was conducted on ln-transformed data.

## Acknowledgments

We sincerely thank Florence Levillain (IPBS, Toulouse) for her training and assistance in the BSL3 facility. We are also grateful to Emmanuelle Näser and Penelope Viana (Génotoul TRI-IPBS platform, Toulouse) for their support with flow cytometry. Additionally, we thank Dr. Pascal Demange (IPBS, Toulouse), Louis Benastre (IPBS, Toulouse), and all members of the Neyrolles lab (IPBS, Toulouse) for their valuable scientific and technical advice. This work was supported by funding from Fondation pour la Recherche Médicale (grant EQU202103012733 to O.N.), Agence Nationale de la Recherche (ANR-21-CE11-0031 BAC-MMEP to O.N. J.M., O.S., and J.-B.S. and ANR-10-INBS-04 FranceBioImaging to J-B.S.), Fondation Bettencourt Schueller (Grant Explore-TB to O.N.), the European Union (Marie Skłodowska-Curie postdoctoral fellowship (MTB-DETOX: 101063199) to P.D.), and NIH grant P30 AI168433 and P30 CA08748 to M.S.G..

## Author contributions

P.D., O.N., C.G., M.S.G., J-B. S. and O.S. designed research; P.D., C.G., Y-M. B., M.F., J.B., A.F., S.C., Y.G., J.M., F.L., L.B., B.V., and J.R. performed research; P.D., O.N., C.G., M.S.G., J-B. S., J.M., and O.S. analyzed data; P.D. and O.N. wrote the paper, with input from all authors. Funding acquisition: O.N., P.D., and J-Y.B.

## Supplementary figures and tables

Supplementary Fig. 1-7 and Supplementary Tables 1-3.

**Movie 1. PacL2 assembles into mobile clusters within the mycobacterial membrane.** Epifluorescence microscopy time-lapse imaging (45-second duration, with one image captured per second) of live *M. smegmatis* expressing the PacL2-mTurquoise protein fusion. Bacteria were cultured in the presence of 10 µM CdSO[. <u>Movie 1</u>.

**Movie 2. Super resolution localization of PacL2 within the mycobacterial membrane.** 3D reconstruction of the PacL2-mEos fusion protein localizations assessed by photoactivated localization microscopy (PALM) in live *M. smegmatis*. Bacteria were cultured in the presence of 10 µM CdSO[. <u>Movie 2</u>.

**Movie 3. The G17L+G20L mutation in PacL2 disrupts the assembly of small mobile clusters within the mycobacterial membrane.** Epifluorescence microscopy time-lapse imaging (45-second duration, with one image captured per second) of live *M. smegmatis* expressing the PacL2^G17L+G20L^-mTurquoise protein fusion. Bacteria were cultured in the presence of 10 µM CdSO[. <u>Movie 3</u>.

**Movie 4. The K9A mutation in PacL2 disrupts the assembly of small mobile clusters within the mycobacterial membrane.** Epifluorescence microscopy time-lapse imaging (45-second duration, with one image captured per second) of live *M. smegmatis* expressing the PacL2^K9A^-mTurquoise protein fusion. Bacteria were cultured in the presence of 10 µM CdSO[. <u>Movie 4</u>.

**Figure S1. Metal sensitivity of *M. smegmatis* strains expressing the PacL/Ctp system under native promoters.** Sensitivity of the indicated *M. smegmatis* strains to specified metals was assessed using a disk diffusion assay. Results are shown as the diameter of the inhibition zone, expressed relative to the empty vector control. Data represent means ± SEM from biological replicates (gray dots). *ctpC* operon: *pacL1-ctpC*. *ctpG* operon: *cmtR-pacL2-ctpG*. *ctpV* operon: *csoR-pacL3-ctpV*. *ctpV* full operon: *csoR-pacL3-ctpV-rv0970*. Statistically significant differences compared to the reference condition are indicated by asterisks (***P < 0.001).

**Figure S2. Metal sensitivity of *M. smegmatis* strains expressing the PacL/Ctp system under an inducible promoter.** Sensitivity of the indicated *M. smegmatis* strains to specified metals was assessed using a disk diffusion assay in (**a**) the absence or (**b**) the presence of the inducer (ATC). Results are shown as the diameter of the inhibition zone. *tet*: ATC-inducible promoter. Data represent means ± SEM from biological replicates (gray dots). Asterisks above the means indicate statistically significant differences compared to the reference strain (empty vector) (***P < 0.001).

**Figure S3. High conservation of amino acid sequences among the three *M. tuberculosis* PacL proteins.** (**a**) Amino acid sequences of *M. tuberculosis* PacL1, PacL2, and PacL3 proteins. (**b**) Sequence alignment of *M. tuberculosis* PacL1, PacL2, and PacL3. Predicted transmembrane domains are shown in blue, AE repeats in red, and putative metal-binding motifs in purple.

**Figure S4. PacL1 Residues bound to Zn or Cd assessed by RMN.** ^1^H-^15^N HSQC overlay spectra of 90 µM ^15^N-labelled SolPacL1 with 0 (black), 0.4 (light blue), 1.0 (purple) and 2.0 (red) equivalent of zinc, spectra (**a**) and (**b**), and cadmium, spectra (**c**) and (**d**). Peak assignment is directly annotated on spectra with the following color code: black (no perturbation), green (perturbed residues in fast exchange) and red with star (perturbed residues in intermediate exchange). We noticed a different binding mode for zinc which shows significant broadening (intermediate chemical exchange) beyond the detection of peaks involved in the metal binding site (H89 to H93) compared to the interaction of SolPacL1 with cadmium which remains mainly in fast exchange, except for H91.

**Figure S5. Modeling of the conserved GXXG motif in PacL2.** (**a**) Sequence conservation logo of 120 bacterial PacL proteins. (**b**) Predicted structural models of the PacL2 transmembrane domain: (left) wild-type (WT) and (right) G17L+G20L mutant.

**Figure S6. Binding of TurboID to the PacL1^ALFA^ proteins does not abolish zinc tolerance.** Anti-ALFA/BirA/RpoB immunoblots from indicated *M. tuberculosis* strains cultivated in the absence or presence of the anhydrotetracycline (ATC) inducer. TurboID refers to the TurboID-nanobody fusion protein under the control of an ATC-inducible (tet-) promoter. *pacL1int-ALFA-ctpC* and *pacL1Cter-ALFA-ctpC* denote the *PacL1* protein carrying an internal or a C-terminal ALFA tag, expressed in operon with *ctpC* under their native promoter. (**b** and **c**) Sensitivity of the indicated *M. smegmatis* strains to ZnSO assessed using a disk diffusion assay in (**b**) absence or (c) presence of 50 nM ATC. Results are shown as the diameter of the inhibition zone, expressed relative to the TurboID + empty vector control. Data represent means ± SEM from biological replicates (gray dots). Statistically significant differences compared to the reference condition are indicated by asterisks (P < 0.001). (**d**) Serial dilutions of *M. tuberculosis* 5 µL cultures spotted on agar plates supplemented or not with 100 µM ZnSO. Three independent biological replicates are shown per condition.

