## Supplementary material for "PacL-organized membrane-associated effluxosomes coordinate multi-metal resistance in *Mycobacterium tuberculosis*": Figure S1

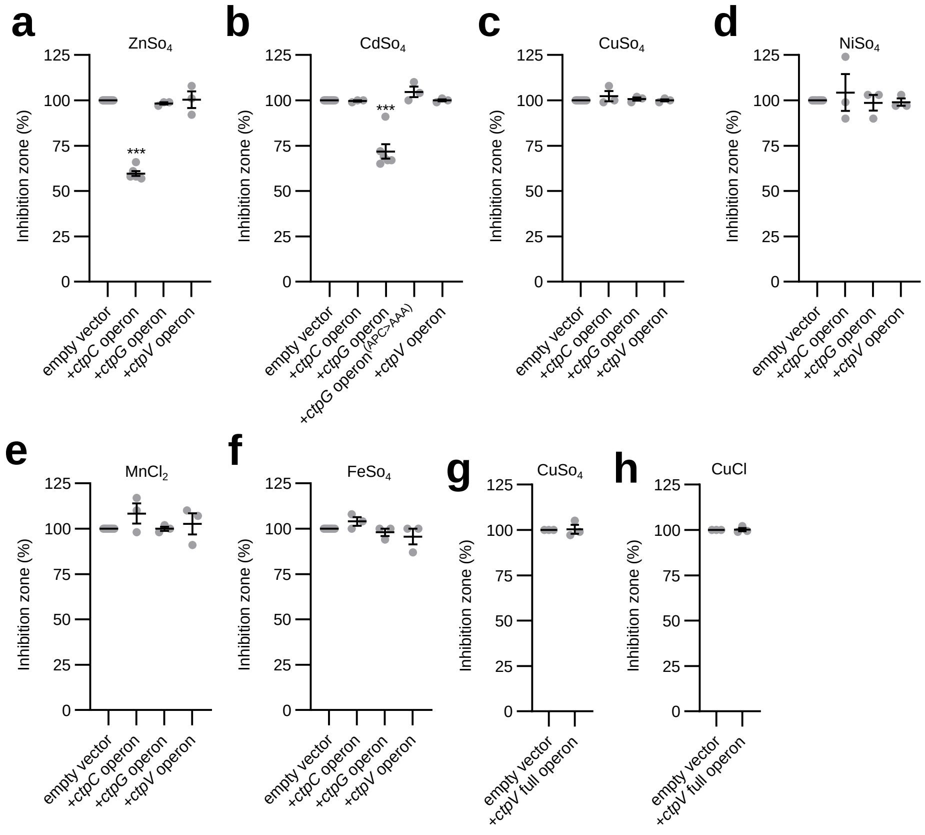


**Figure S1. Metal sensitivity of *M. smegmatis* strains expressing the PacL/Ctp system under native promoters.** Sensitivity of the indicated *M. smegmatis* strains to specified metals was assessed using a disk diffusion assay. Results are shown as the diameter of the inhibition zone, expressed relative to the empty vector control. Data represent means ± SEM from biological replicates (gray dots). *ctpC* operon: *pacL1-ctpC*. *ctpG* operon: *cmtR-pacL2-ctpG*. *ctpV* operon: *csoR-pacL3-ctpV*. *ctpV* full operon: *csoR-pacL3-ctpV-rv0970*. Statistically significant differences compared to the reference condition are indicated by asterisks (***P < 0.001).
