## Supplementary material for "PacL-organized membrane-associated effluxosomes coordinate multi-metal resistance in *Mycobacterium tuberculosis*": Figure S2

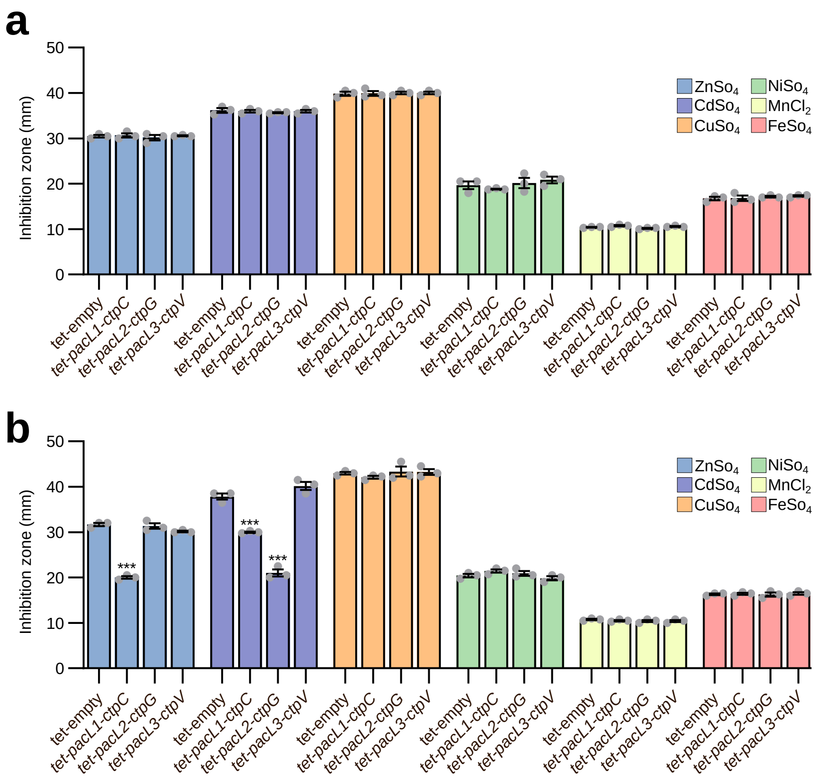


**Figure S2. Metal sensitivity of *M. smegmatis* strains expressing the PacL/Ctp system under an inducible promoter.** Sensitivity of the indicated *M. smegmatis* strains to specified metals was assessed using a disk diffusion assay in (**a**) the absence or (**b**) the presence of the inducer (ATC). Results are shown as the diameter of the inhibition zone. *tet*: ATC-inducible promoter. Data represent means ± SEM from biological replicates (gray dots). Asterisks above the means indicate statistically significant differences compared to the reference strain (empty vector) (***P < 0.001).
