## Supplementary material for "PacL-organized membrane-associated effluxosomes coordinate multi-metal resistance in *Mycobacterium tuberculosis*": Figure S3

**a**

```
>Rv3269(PacL1)
MAIQVFLAKATTTVITGLAGVTAYEILKKAAAKAPLRQTAVSAAALGLRGTRKAEEAAESARLKVADVMA
EARERIGEESPTPAISDLHDHDH

>Rv1993c(PacL2)
MVTHELLVKAAGAVLTGLVGVSAYETLRKALGTAPIRRASVTVMEWGLRGTRRAEAAAESARLTVADVVA
EARGRIGEEAPLPAGARVDE

>Rv0968(PacL3)
MVWHGFLAKAVPTVVTGAVGVAAYEALRKMVVKAPLRAATVSVAAWGIRLAREAERKAGESAEQARLMFA
DVLAEASERAGEEVPPLAVAGSDDGHDH
```

**b**

```
PacL1 MAIQVFLAKATTTVITGLAGVTAYEILKKAAAKAPLRQTAVSAAALGLRG
PacL2 VVTHELLVKAAGAVLTGLVGVSAYETLRKALGTAPIRRASVTVMEWGLRG
PacL3 MVWHGFLAKAVPTVVTGAVGVAAYEALRKMVVKAPLRAATVSVAAWGIRL
      :. : :*.**. :*:** .**:*** *:*   .**:* ::*:.   *:*

PacL1 TRKAE----EAAESARLKVADVMAEARERIGEESPTPAISDLHD-HDH
PacL2 TRRAE----AAAESARLTVADVVAEARGRIGEEAPLPAGARVDE----
PacL3 AREAERKAGESAEQARLMFADVLAEASERAGEEVPPLAVAGSDDGHDH
      :*.**     :**.*** .***:***  * *** *  * :  .:
```

**Figure S3. High conservation of amino acid sequences among the three *M. tuberculosis* PacL proteins.** (**a**) Amino acid sequences of *M. tuberculosis* PacL1, PacL2, and PacL3 proteins. (**b**) Sequence alignment of *M. tuberculosis* PacL1, PacL2, and PacL3. Predicted transmembrane domains are shown in blue, AE repeats in red, and putative metal-binding motifs in purple.
