## Supplementary material for "PacL-organized membrane-associated effluxosomes coordinate multi-metal resistance in *Mycobacterium tuberculosis*": Figure S4

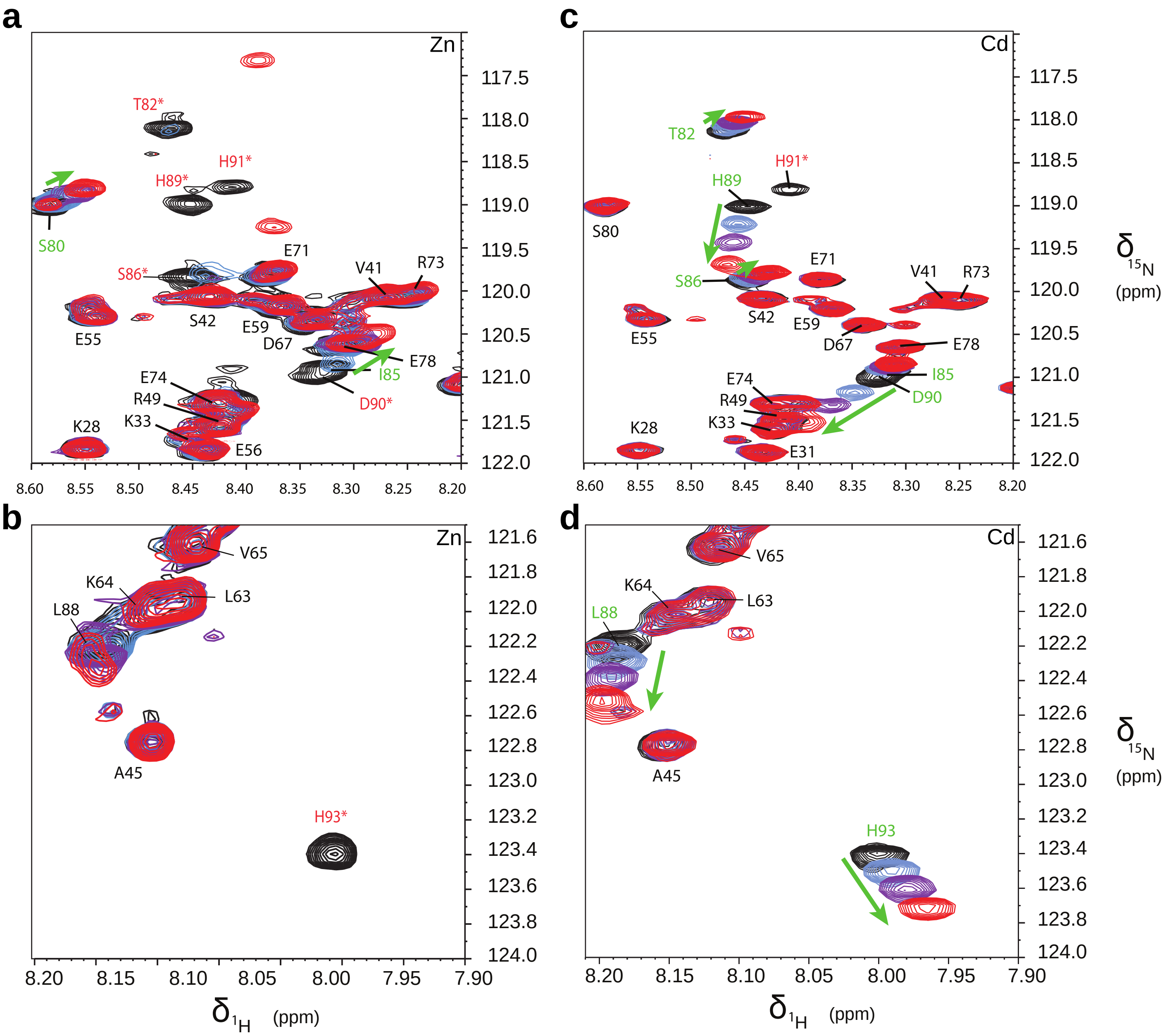


**Figure S4. PacL1 Residues bound to Zn or Cd assessed by RMN.** $^{1}$H-$^{15}$N HSQC overlay spectra of 90 µM $^{15}$N-labelled SolPacL1 with 0 (black), 0.4 (light blue), 1.0 (purple) and 2.0 (red) equivalent of zinc, spectra (**a**) and (**b**), and cadmium, spectra (**c**) and (**d**). Peak assignment is directly annotated on spectra with the following color code: black (no perturbation), green (perturbed residues in fast exchange) and red with star (perturbed residues in intermediate exchange). We noticed a different binding mode for zinc which shows significant broadening (intermediate chemical exchange) beyond the detection of peaks involved in the metal binding site (H89 to H93) compared to the interaction of SolPacL1 with cadmium which remains mainly in fast exchange, except for H91.
