## Supplementary material for "PacL-organized membrane-associated effluxosomes coordinate multi-metal resistance in *Mycobacterium tuberculosis*": Figure S5

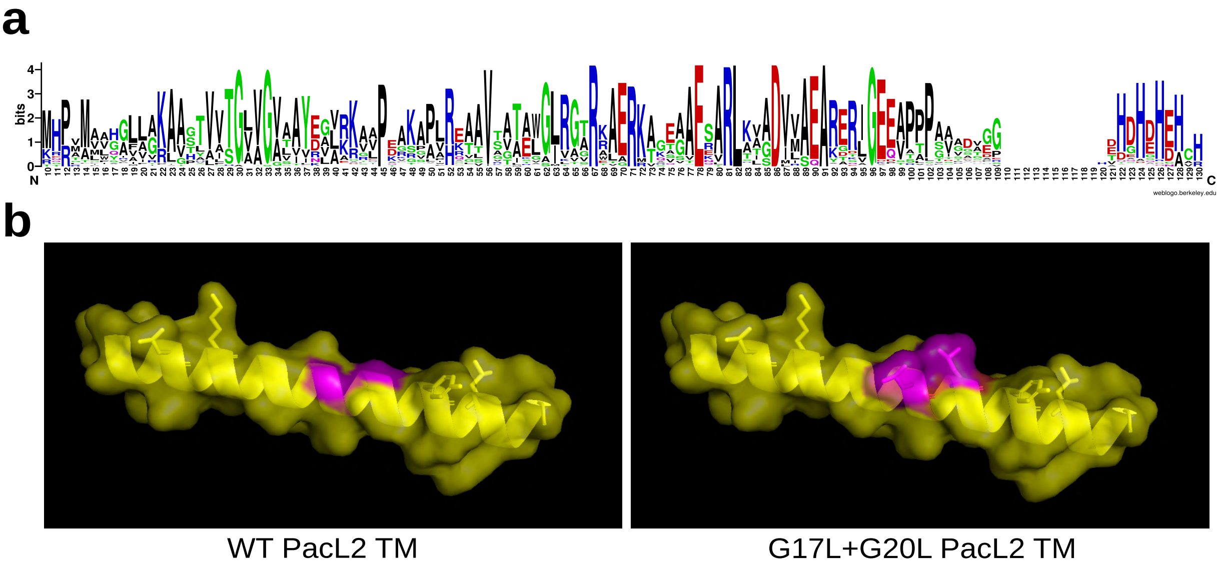


**Figure S5. Modeling of the conserved GXXG motif in PacL2.** (**a**) Sequence conservation logo of 120 bacterial PacL proteins. (**b**) Predicted structural models of the PacL2 transmembrane domain: (left) wild-type (WT) and (right) G17L+G20L mutant.
