## Supplementary material for "PacL-organized membrane-associated effluxosomes coordinate multi-metal resistance in *Mycobacterium tuberculosis*": Figure S6

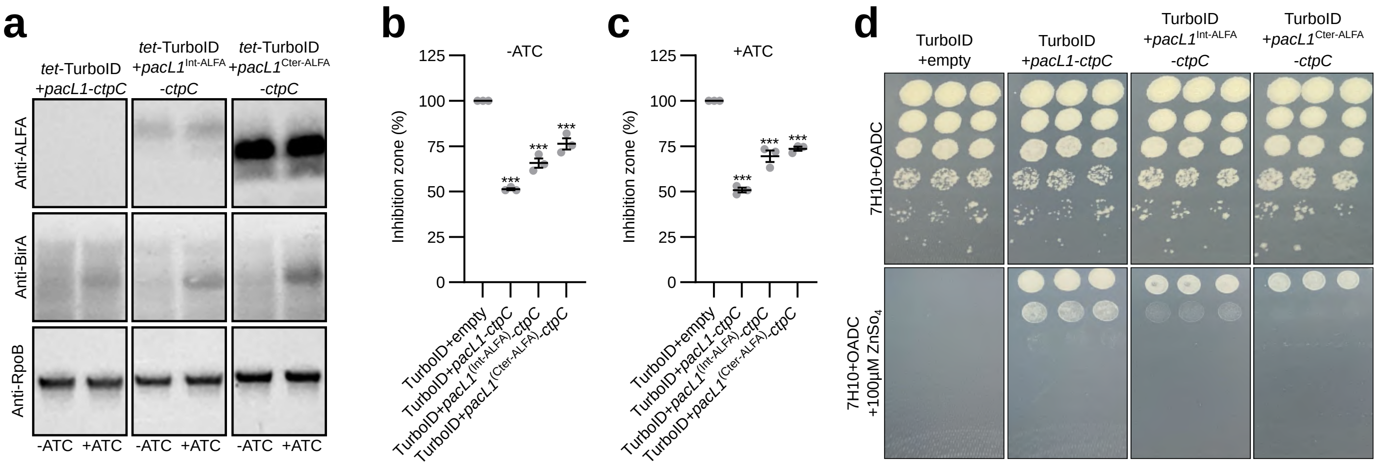


**Figure S6. Binding of TurboID to the PacL1$^{ALFA}$ proteins does not abolish zinc tolerance.** Anti-ALFA/BirA/RpoB immunoblots from indicated *M. tuberculosis* strains cultivated in the absence or presence of the anhydrotetracycline (ATC) inducer. TurboID refers to the TurboID-nanobody fusion protein under the control of an ATC-inducible (tet-) promoter. *pacL1int-ALFA-ctpC* and *pacL1Cter-ALFA-ctpC* denote the *PacL1* protein carrying an internal or a C-terminal ALFA tag, expressed in operon with *ctpC* under their native promoter. (**b** and **c**) Sensitivity of the indicated *M. smegmatis* strains to $ZnSO_4$ assessed using a disk diffusion assay in (**b**) absence or (c) presence of 50 nM ATC. Results are shown as the diameter of the inhibition zone, expressed relative to the TurboID + empty vector control. Data represent means ± SEM from biological replicates (gray dots). Statistically significant differences compared to the reference condition are indicated by asterisks ($P < 0.001$). (**d**) Serial dilutions of *M. tuberculosis* 5 µL cultures spotted on agar plates supplemented or not with 100 µM $ZnSO_4$. Three independent biological replicates are shown per condition.
